# Category-selective human brain processes elicited in fast periodic visual stimulation streams are immune to temporal predictability

**DOI:** 10.1101/117135

**Authors:** Genevieve L. Quek, Bruno Rossion

## Abstract

Recording direct neural activity when periodically inserting exemplars of a particular category in a rapid visual stream of other objects offers an objective and efficient way to quantify perceptual categorization and characterize its spatiotemporal dynamics. However, since periodicity entails predictability, perceptual categorization processes identified within this framework may be partly generated or modulated by temporal expectations. Here we present a stringent test of the hypothesis that temporal predictability generates or modulates category-selective neural processes. In Experiment 1, we compare neurophysiological responses to periodic and nonperiodic (i.e., unpredictable) variable face stimuli in a fast (12 Hz) visual stream of nonface objects. In Experiment 2, we assess potential responses to rare (10%) omissions of periodic face events (i.e., violations of periodicity) in the same fast visual stream. Overall, our observations indicate that long duration category(face)-selective processes elicited in a fast periodic stream of visual objects are immune to temporal predictability. These observations do not support a predictive coding framework interpretation of category-change detection in the human brain and have important implications for understanding human perceptual categorization in a rapidly changing (i.e., dynamic) visual scene.

---

The human brain organizes visual information in its environment both rapidly and effortlessly. With just a single glance, we can tell almost instantly that a roundish object in our field of view is a face - not a flower, a tennis ball, a clock, or any other type of object. This ability to rapidly group stimuli into meaningful categories - known as perceptual categorization - is surely one of the most fundamental high-level brain functions, serving as the foundation for memory, learning, language, affective processing, decision making, and action execution.

In the visual domain, a powerful way to shed light on perceptual categorization processes is to combine visual periodicity with direct recording of neural activity, for instance using electroencephalography (EEG). By embedding members of a specific category at a strict periodic rate within a dynamic visual stream of items that do *not* belong to that category, it is possible to project the perceptual categorization processes of interest to a specified frequency in the EEG spectrum. At a rapid (and quasi-continuous) rate, this approach can isolate category-selective visual processes without post-hoc subtraction, in a manner that is both objective and highly efficient. For example, Lochy, Van Belle, and Rossion (2015) investigated lexical categorization processes by presenting participants with a stream of non-word items a rate of exactly 10 Hz (i.e., 10 non-words per second), with a word stimulus embedded as every fifth item. Three minutes of this stimulation elicited an electrophysiological response at the exact frequency of image presentation (i.e., 10 Hz), but more importantly, a robust response at the exact periodicity of the word items embedded in non-word sequence (i.e., 10 Hz/5 items = 2 Hz), even in the absence of an overt lexical task. The authors interpreted this 2 Hz signal to be a differential response to words compared to non-words, as it could only have arisen if the response evoked by words *differed* from that evoked by non-words (see also Lochy et al. 2016). The same periodicity-based approach has also been used to examine human adults and infants’ perceptual categorization of faces and natural object images (e.g., Figure 1; de Heering and Rossion 2015; Rossion et al. 2015; Jacques et al. 2016; Retter and Rossion 2016). For example, Retter and Rossion (2016) presented participants with a dynamic stream of object images at a rate of 12.5 Hz (i.e., 80 ms per image), inserting face images into the sequence every three, five, seven, nine, or 11 stimuli. In addition to finding a strong response at the image presentation frequency of 12.5 Hz, they also observed a robust category-selective response at each of the defined face periodicities (e.g., a face every seven images gives a response at exactly 12.5 Hz/7 = 1.79 Hz). Given the very rapid image presentation rate (each image was replaced after just 80 ms), and the use of a completely orthogonal task, the authors argued that this category-selective response reflected automatic categorization of faces vs. objects at the perceptual rather than decisional level. This conclusion is supported by the application of this approach to intracerebral recordings in a large group of human patients, identifying and quantifying the face-selective responses primarily in localized regions of the right ventral occipito-temporal cortex (Jonas and Rossion 2016).

**Figure 1.**
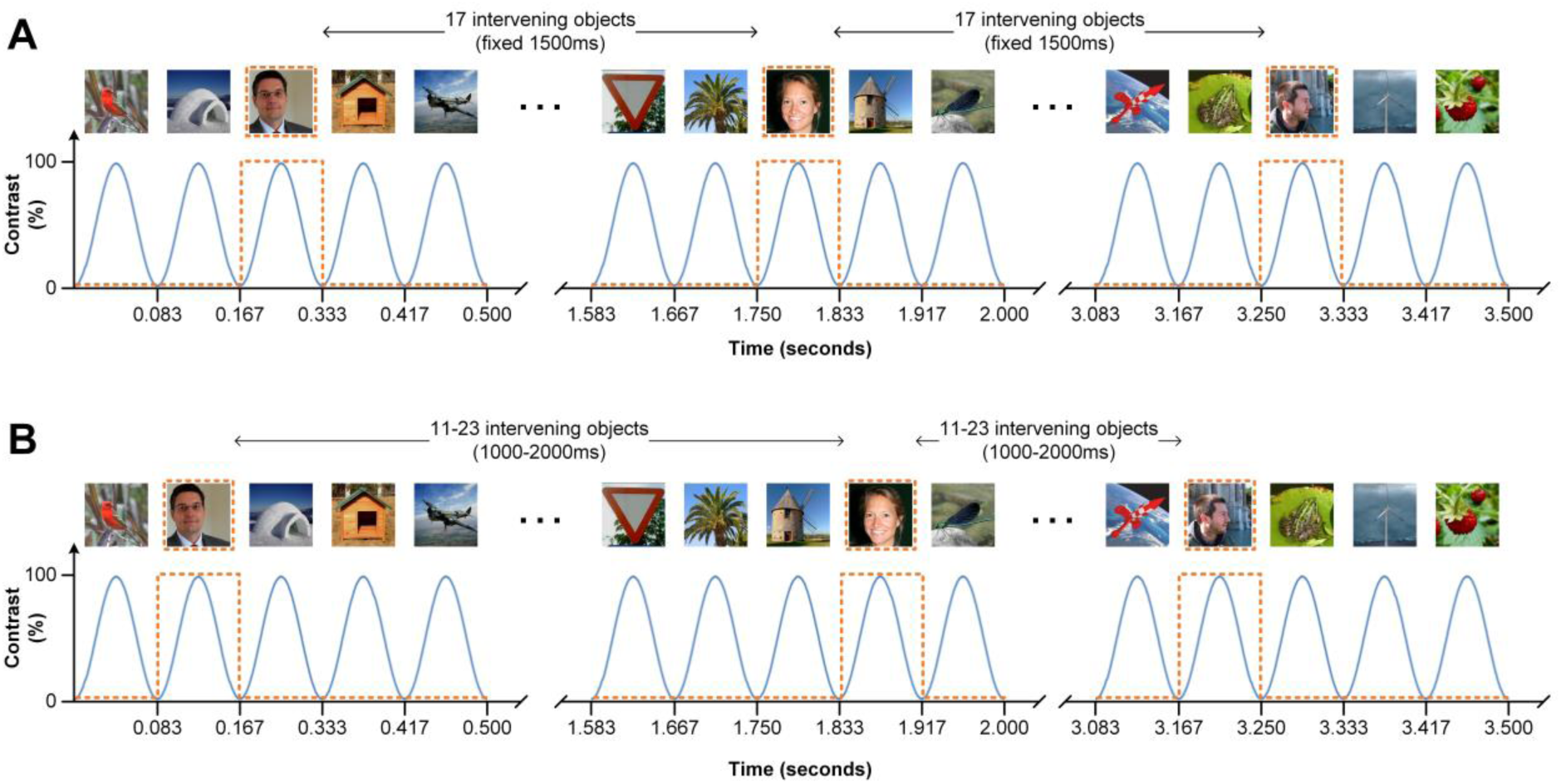
Experiment 1 sequence design. We presented participants with 12 images per second (12 Hz), sinusoidally modulating the contrast of each from 0-100-0% (blue solid lines). In both conditions, the 90 s stimulation sequence contained 60 natural face images and numerous natural object images (e.g., vehicles, animals, buildings, trees, etc.). (A) In the periodic condition, faces were spaced at regular intervals during the stimulation sequence, appearing every 18 stimuli (orange dashed lines). (B) In the nonperiodic condition, faces were spaced at irregular intervals, appearing anywhere after 11 to 23 object images (orange dashed lines). Participants did not respond to the faces during the sequence, but instead monitored a central fixation cross overlaid on the images for changes of color.

Since the category-selective response in dynamic visual streams depends on *i)* the critical stimuli and temporally surrounding distractors evoking different responses, and *ii)* measuring a similar evoked response each time an exemplar of the critical category appears, this approach (known as fast periodic visual stimulation, or FPVS) provides a way to rapidly and objectively quantify processes related to both visual discrimination and visual generalization. An important and outstanding theoretical issue, however, is whether the category-selective signal yielded by periodicity is generated in part by temporal expectation. That is, since the critical category exemplars always appear at periodic intervals in FPVS, and since the entire sequence is itself a rhythmic stimulation (Jones 1976), participants in these tasks could conceivably form reliable expectations (either explicit or implicit) about exactly *when* critical stimuli will appear (McAuley and Jones 2003). Indeed, a number of studies have shown that regular (“rhythmic”) stimulation can induce strong temporal expectations, thereby facilitating sensory processing of stimuli both in the auditory (e.g., Morillon et al. 2016) and visual domains (Mathewson et al. 2010; Rohenkohl et al. 2012; Cravo et al. 2013; Breska and Deouell 2014). Generally, these effects are expressed in terms of greater encoding precision, higher perceptual sensitivity and decreased response times in behavioral tasks (Rajendran and Teki 2016). Moreover, behavioral studies employing rapid serial visual presentation (RSVP; Potter and Levy 1969), a stimulation that is similar in kind to FPVS, have shown that identification accuracy for targets embedded in these streams improves as a function of number of distractors before target onset, suggesting that temporal expectation is “tuned” over the course of the RSVP sequence itself (Ariga and Yokosawa 2008).

If the rhythmicity of the FPVS approach (e.g., images appearing at a defined periodic rate), combined with the temporal predictability of critical category exemplars (e.g., a face after every 9 object images), does indeed elicit temporal expectations in participants, then the category-selective response it yields may not solely reflect processes related to perceptual categorization, but may be generated in part by temporal expectation. As a case in point, the category-selective response for faces embedded in a stream of objects is known to be comprised of several components starting at ~100ms and lasting up to ~500ms after face onset (Rossion et al. 2015; Jacques et al. 2016; Retter and Rossion 2016). Given that behavioral detection of a face can be achieved within 150-300ms following stimulus onset (Rousselet et al. 2003; Crouzet et al. 2010; Crouzet and Thorpe 2011), it could well be the case that the later aspects of this neural response may be generated or at least largely modulated by temporal expectation about the appearance of faces in the visual stream. Indeed, faces are perhaps the ideal stimulus with which to test the role of temporal expectation in driving the category-selective response in periodicity-based designs, since observers can easily form a visual template for this category based on the cardinal arrangement of facial features. Moreover, face perception is known to be highly sensitive to top-down effects, as evidenced by phenomena like the hollow-face illusion (Gregory 1970) and face recognition of 2-tone pictures ("Mooney faces"; Mooney 1957; Cavanagh and Leclerc 1989; Moore and Cavanagh 1998).

Here we present two EEG experiments that shed light on the important and unresolved issue of whether temporal predictability modulates implicit perceptual categorization in the context of dynamic visual stimulation. In Experiment 1, we compare the category-selective response elicited by FPVS sequences in which the critical face stimuli are either temporally predictable (embedded at periodic intervals) or unpredictable (embedded at nonperiodic intervals). In Experiment 2, we examine the neural response to a violation of temporal predictability in an FPVS sequence, by introducing rare random omissions of periodic (i.e., expected) faces after a periodic of entrainment.

## 1. Experiment 1

Our goal in Experiment 1 was to dissociate the relative contribution of two factors that are typically conflated in FPVS designs - periodicity and temporal predictability. To this end, we embedded face stimuli at either fixed or random intervals in a sequence of variable object images. After verifying this manipulation in the frequency-domain, we employed a novel ‘False-Sequencing’ approach to impose a new face periodicity of exactly 1 Hz on both the predictable (i.e., periodic) and unpredictable (i.e., nonperiodic) conditions. In this analysis, face periodicity is held constant (always 1 Hz) while temporal predictability is allowed to vary. If temporal predictability confers a processing advantage on the critical face stimuli in FPVS sequences, then the category-selective response elicited by temporally predictable and unpredictable faces should differ either quantitatively or qualitatively. Under a predictive coding framework, we might expect this difference to take the form of an attenuated response to periodic faces as compared to nonperiodic faces, since sensory input corresponding to the former stimulus is consistent with current high level expectations and thus effectively redundant (Rao and Ballard 1999; Friston 2005; Alink et al. 2010; Kok, Jehee, et al. 2012). On the other hand, if temporal expectation sharpens sensory representation by suppressing neural responses that are inconsistent with current expectations (Lee and Mumford 2003), then periodic faces might be expected to elicit a stronger category-selective response than nonperiodic faces, since noise is reduced in the former case.

## 2. Experiment 1 Methods

### 2.1. Participants

Twenty persons (nine females) aged between 18-26 years took part in this study in exchange for payment. All participants were right-handed, with normal or corrected-to-normal vision, and no history of neurological illness. In accordance with the University of Louvain Biomedical Ethics Committee guidelines, we obtained written informed consent from all participants prior to testing.

### 2.2. Stimuli

Stimuli were 200 color images of various non-face objects (e.g., animals, plants, manmade structures/objects) and 100 color images of human faces. We resized each image to 256x256 pixels and equalized their mean luminance values in MATLAB (R2012b). As in our previous studies that used a different but comparable stimulus set (e.g., Rossion et al. 2015), we deliberately did not segment faces/objects from their original naturalistic backgrounds, such that the viewpoint, lighting, and composition images varied widely across the full stimulus set (see Figure 1 for examples). During the stimulation sequence, each image appeared on a uniform grey background and subtended 8.721° of visual angle. A small central fixation cross (subtending 0.788° of visual angle) overlaid the images throughout the sequence.

### 2.3. Procedure

Participants sat in a darkened room and viewed a computer monitor at a distance of 80cm. Each stimulation sequence consisted of a two second pre-stimulation period that contained only the central fixation cross, a two second fade-in, a 90 second sequence of rapid images presented at a periodic rate of exactly 12 Hz (12 images per second), and a three second fade-out. We achieved this 12 Hz presentation rate using customized software programmed in Java that sinusoidally modulated the contrast of each image from 0% to 100% to 0% over a period of 83.33 ms (see Figure 1; Movie 1). Since the stimuli are still visible at low contrast, sinusoidal contrast modulation ensured that the stimulation was near-continuous, such that participants did not perceive any interruption between images (see Retter and Rossion 2016, for a comparison of sinusoid and squarewave stimulation at 12.5 Hz). We instructed participants to fix their gaze on the central cross overlaid on the images, and to press the SPACE key whenever this cross changed color from black to red (200ms change duration; 12 changes in each stimulation sequence; minimum 2 seconds (s) between consecutive changes). In the periodic condition, sequences consisted of randomly selected non-face images with a face image embedded as every 18^th^ stimulus (Figure 1A). Presenting faces at fixed intervals during the sequence should elicit two periodic EEG responses: one at 12 Hz, reflecting visual processing common to both face and non-face stimuli (referred to as the common response), and one at 12 Hz/18 (i.e., 0.67 Hz), reflecting the differential response between faces and the other images (referred to here as the category-selective or face-selective response, Retter & Rossion, 2016). Nonperiodic sequences were identical to periodic sequences, save that face images appeared at *irregular* intervals during the rapid sequence (11-23 non-face images between the face images, see Figure 1B). Here only the common response at 12 Hz should be evident in the EEG spectrum, since there is no second periodicity embedded in the sequence. Since previous studies have observed significant category-selective responses in most participants after just 2-4 stimulation sequences, here participants viewed four periodic and four nonperiodic stimulation sequences in a pseudorandom order, each containing exactly 60 face image presentations.

### 2.4. EEG Acquisition

We acquired high-density 128-channel EEG during sequence viewing using an ActiveTwo Biosemi system with a 512 Hz sampling rate (http://www.biosemi.com/). We monitored eye-movements via four external electrodes placed at the outer canthi of the eyes, and above and below the right eye. The electrode offset criterion cut-off was 20μV

### 2.5. Analysis

#### 2.5.1. Behavioral Analysis

We calculated the accuracy and response time (RT) for every fixation cross color change during the stimulation sequence, removing values that occurred < 250ms or > 1500ms following change onset.

#### 2.5.2. EEG Analysis

We analyzed EEG data using the open source toolbox Letswave5 (http://nocions.webnode.com/letswave) running over MATLAB R2012b (MathWorks, USA), and R (https://www.r-proiect.org/). We note that all EEG waveform analysis procedures described here have also been extensively documented in recent publications (Liu-Shuang et al. 2014; Rossion 2014; Rossion et al. 2015; Jacques et al. 2016; Retter and Rossion 2016).

##### 2.5.2.1. EEG pre-processing

We excluded data from two male participants due to excessive artefact on multiple channels in the EEG trace. Before removing these participants, we verified that both showed a significant face-selective response in the EEG spectrum. For each of the remaining 18 participants, we removed slow voltage changes of non-neural origin and irrelevant EMG artefact by applying a 4^th^ order Butterworth band-pass filter (0.05-100 Hz) to the raw EEG trace (Luck, 2005). We then removed AC electrical noise at 50 Hz, 100 Hz, and 150 Hz with an FFT multi-notch filter (width = 0.5 Hz), and then segmented 99 s epochs relative to the starting trigger of each stimulation sequence (-2 s to 97 s). To remove blink artefact, we applied an independent components analysis (ICA) with a square mixing matrix (Hyvarinen and Oja 2000) and removed a single component identified by visual inspection of the waveform and corresponding topography. Artefact-prone channels with deflections greater than 100 µV in at least two trials were replaced using linear interpolation of neighboring clean channels (less than 5% of channels per participant). Since we recorded high-density EEG here, we re-referenced each channel’s signal using the mean of all 128 scalp channels (rather than single reference electrode), and relabeled each to match the standard 10-20 system (for additional detail see Rossion et al. 2015).

##### 2.5.2.2. Frequency-domain analysis

Following pre-processing, we segmented an epoch for each sequence containing an integer number of cycles of the face presentation frequency (0.67 Hz), resulting in an 88.51 second epoch containing exactly 59 face presentation cycles for each stimulation sequence. This ensured that a single frequency bin would be centered on the category-selective frequency of interest (0.67 Hz) in the EEG spectrum. We averaged the four epochs for each condition before applying a Fast Fourier Transformation (FFT) to extract a normalized amplitude spectrum for each channel ranging between 0 and 256 Hz. The frequency resolution of these spectra was very high, as determined by the inverse of the sequence duration (i.e., 1/88.5 = 0.0113 Hz). To identify significant response signal at the relevant stimulation frequencies (i.e., 12 Hz, 0.67 Hz, and harmonics), we computed a *z*-score at each discrete frequency bin (e.g., Liu-Shuang et al. 2014; Rossion 2014; Rossion et al. 2015; Jacques et al. 2016; Retter and Rossion 2016) using the amplitude spectra pooled across all participants, conditions, and scalp channels. Here we specified a noise range comprised of the 20 frequency bins neighboring the frequency of interest, excluding the immediately adjacent bins and the local maximum and minimum amplitude bins (e.g., Rossion and Boremanse 2011; Rossion et al. 2012; Retter and Rossion 2016). We subtracted the mean amplitude of this noise range from the amplitude at the frequncy of interest, and divided the result by the standard deviation of amplitudes in the noise range. The advantage of determining statistical significance this way is that unlike *t*-tests (which rely in inter-individual variance), z scores can computed either at the group level or for each participant individually (for an extended discussion of detecting significant signal at a specified frequency, see Appendix 2 of Norcia et al. 2015). As in our previous studies (e.g., Jacques et al. 2016), we considered response signals with a z-score greater than 3.1 to be significant (*p* < .001, one tailed, i.e., signal > noise), and selected only these frequencies for further analysis.

To take into account the variation in noise across the EEG spectrum, we performed a local baseline-subtraction on the raw amplitudes for each condition using the same baseline range as for z-score calculation (see above). We then summed the baseline-corrected amplitudes across all relevant harmonic frequencies (Jacques et al. 2016; Retter and Rossion 2016). As in previous studies, we performed this quantification of the category-selective response two ways. First, we compared the response in each condition at the global level by pooling the information across all scalp channels. Second, we compared the response in functional regions-of-interest (ROIs) compatible with the stable bilateral occipito-temporal pattern typically elicited by FPVS paradigms (Rossion et al. 2015; Jacques et al. 2016; Retter and Rossion 2016). We defined these ROIs by averaging the four channels in each hemisphere with the greatest summed-harmonic response averaged across-conditions, resulting in a left occipito-temporal ROI (averaged over channels P9, PO7, PO9, and PO11) and a right occipito-temporal ROI (averaged over channels P10, PO8, PO10, and PO12). The central occipito-parietal ROI was the average of the four channels with the largest common response at 12 Hz (averaged over channels Oz, POOz, PO4h, and POO6).

##### 2.5.2.3 False-sequencing analysis

To dissociate the contribution of periodicity and temporal predictability to category-selective response, we used a novel approach we term ‘false-sequencing’ here. For each stimulation sequence in each condition, we segmented a 1000ms epoch around each face onset (-500ms to 500ms), resulting in ~60 segments per stimulation sequence. We sequenced these 1000ms segments by aligning the first sample of each epoch with the last sample of the preceding epoch, producing a new 60 second continuous EEG trace. To compensate for introduced drift in this newly created ‘false-sequence’, we subtracted the mean voltage on each channel and linearly de-trended the data. We then cropped each false-sequence to be exactly 55 s (corresponding to an integer number of number of cycles of 0.67 Hz), and averaged these together by condition. Remaining analyses for the false-sequence data were as described for the frequency-domain analysis above. Additionally, we used the ‘BayesFactor’ package in R (http://bavesfactorpcl.r-forge.r-proiect.org/) to calculate a Bayes factor for each possible model of the false-sequence data containing the factors *Periodicity* and *ROI.*

##### 2.5.2.4. Time-domain analysis

To compare the response to periodic and nonperiodic faces in the time-domain, we applied a standard low-pass filter to the raw EEG trace using a 30 Hz cut-off (4^th^ order Butterworth filter) before cropping each stimulation sequence to an integer number of cycles of the image presentation rate (i.e., 12 Hz). To isolate the differential response to faces (as opposed to objects), we applied a multi-notch filter (0.05 Hz width) at 12 Hz, 24 Hz, and 36 Hz to selectively remove signal corresponding to the image presentation rate. For each face occurrence in the resulting data, we segmented a 1000 ms epoch around face onset (-200ms to 800ms). We then averaged the ~240 epochs for each condition, and applied a standard baseline-correction procedure by subtracting the response amplitude averaged over 167 ms preceding face onset from the waveform. In each of our three ROIs, we compared the conditional mean waveforms by conducting a paired *t*-test at each of the 513 time points. We inspected these obtained *t*-values relative to two significance criteria (*p* < .01 uncorrected, and *p* < .05, corrected for multiple comparisons). We corrected for multiple comparisons using a permutation procedure described by Blair and Karniski (1993; see also Finkbeiner and Friedman 2011) which maintains experiment-wise error in ERP analysis. The steps for this procedure were as follows. First we generated 100,000 permutations of our data by systematically shuffling the periodic/nonperiodic labels for each participant’s conditional mean datasets (consisting of 513 time points). Assuming the null hypothesis is true (i.e., there is no difference between periodic and nonperiodic waveforms), then the assignment of condition labels will be arbitrary, such that each instance of permutated data is just as likely to have occurred as our actual obtained data. In this way, the permuted data arrangements represent 100,000 possible outcomes of our experiment. For each permutation, we conduct a paired t-test at every time point (*n* = 513, for a total of 51.3 million comparisons) and keep aside the maximum *t*-value (*t*_|max|_) given by that permutation. By rank ordering the *t*_|max|_ values obtained across all permutations, we form a reference distribution against which the obtained test statistic at each time point can be compared. To maintain experiment-wise α at 0.05, we set our critical *t* to be the value in the *t*_|max|_ reference distribution that cuts off 0.025 of each tail (see Blair and Karniski, 1993 for a more detailed explanation of the rationale behind this procedure).

We assessed the similarity/difference of the time-domain response to periodic and nonperiodic faces in a second way using multivariate pattern classification (MVPA). Unlike the permutation testing described above, MVPA does not rely on single channel waveform data (i.e., a single averaged channel for each ROI), but considers fine differences in the pattern of activity elicited by periodic and nonperiodic faces across all channels simultaneously. We used an MPVA procedure for spatiotemporal decoding in EEG introduced by Bode and colleagues (The Decision Decoding Toolbox, available at http://ddtbox.github.io/DDTBOX/; Bode and Stahl 2014; Bode et al. 2016). Using the low-pass filtered data in which the image presentation frequency of 12 Hz was preserved (i.e., not yet filtered out), we extracted epochs corresponding to periodic faces and nonperiodic faces (- 166.66ms to 500ms) for each participant. We used non-overlapping spatiotemporal analysis time-windows of 20 ms, each containing 10 data points for each of the 128 channels. This time-window moved forward by 20 ms increments to cover the entire 666.66 ms epoch (Bai et al. 2007; Das et al. 2010; Blankertz et al. 2011; Bode et al. 2012; Bode and Stahl 2014). At each time-window, we transformed the data into a spatiotemporal pattern vector with its periodicity condition. We then trained an SVM classifier on the periodic face and nonperiodic face vectors using LIBSVM (Chang and Lin 2011) using 90% of the data, and tested it on the remaining 10%. We repeated this classification process using a 10-fold cross-validation procedure which took different, non-overlapping chunks of test data and train data on each pass. To further guard against any drawing biases that could influence classification, we repeated the ten cross-validation steps five times over, each time with a newly drawn 10% test portion of the data. We calculated classification accuracy at each time-window by averaging across these 50 analyses, and compared each resulting value to an empirical chance distribution corresponding to that time-window. This reference distribution was generated using the exact same classification procedure with randomly shuffled labels for each iteration (i.e., each face epoch randomly labelled as either periodic or nonperiodic). Collated across participants, classification accuracies for the permutated data form a reference distribution at each time point, the mean of which we compared to the real classification accuracy using a paired *t*-test (34 time points, significance criterion = *p* < .05, Bonferroni corrected). This method is considered stricter than testing against theoretical chance accuracy of 50% (Bode and Stahl 2014; Bode et al. 2016).

In a complementary analysis, we used MVPA to ask whether the temporal unfolding of activity across the scalp elicited by a periodic face would *generalize* well to the activity elicited by a nonperiodic face. Again using non-notch filtered data, here we segmented an equal number of face and object instances for each periodicity condition and participant (- 166.66ms to 500ms). We ensured an equal number of both stimulus types by taking only those object instances appearing exactly 6 cycles before each face. Using the same classification parameters as described above, we performed i) a <u>within-condition</u> decoding analysis of faces vs. objects by (classifier trained using 90% of face and object epochs from the *periodic* condition, and tested it on the remaining 10% of epochs from that same condition), and ii) a <u>cross-condition</u> decoding analysis (classifier on 90% of the *periodic* data, and tested it on 10% of the *nonperiodic* data). The rationale for cross-condition decoding is as follows - since the classifier is trained to distinguish neural activity evoked by an object from that evoked by a temporally predictable face, if it can perform at above-chance level in classifying neural activity elicited by objects and temporally unpredictable faces, then the response evoked by faces under predictable and unpredictable conditions must be similar. Alternatively, if the classifier trained to distinguish periodic face and object responses cannot generalize to nonperiodic face and object data, then we may conclude that response evoked by temporally regular and irregular faces is qualitatively different. In a final step, we compared the decoding accuracy for these two analyses at each time point using paired *t*-tests (34 time points, significance criterion: *p* < .05, False Discovery Rate corrected). If cross-condition decoding (i.e., generalizing from periodic data to nonperiodic data) is significantly worse than within-condition decoding (i.e., generalizing from periodic data to periodic data), we may conclude that face periodicity changes the nature of the face response in FPVS paradigms.

## 3. Experiment 1 Results

### 3.1. Behavior

Accuracy for the fixation cross color change task was uniformly high in both conditions, with no significant difference in percent correct between periodic trials (*M* = 95.95%, *SD* = 8.44%) and nonperiodic trials (*M* = 96.41%, *SD* = 6.46%), *t*(17) = .403, *p* = .692, *d* = 0.09. There was also no evidence that mean response time (RT) differed between the periodic (*M* = 483ms, SD = 51ms) and nonperiodic conditions (*M* = 485ms, *SD* = 53ms), *t*(17) = .648, *p* = .526, *d* = 0.15.

### 3.2. Frequency-Domain Results

#### 3.2.1. Response at the category-selective frequency (0.67 Hz).

To avoid condition or channel related biases, we determined the range of frequencies for quantification of the category-selective response by inspecting the amplitude spectrum averaged across all conditions and all channels. There was a highly significant response (i.e., *p* < .001, one tailed, indicating signal > noise) at each the first 13 consecutive harmonics associated with the category-selective frequency (i.e., 0.67 Hz). When split by condition, the global scalp response at all 13 of these harmonics was significant in the periodic condition, but not the nonperiodic condition. Accordingly, the sum of baseline-corrected amplitude across these 13 harmonics was much larger in the periodic condition (*M* = 1.162 μV, *SD* = .466 µV) compared to the nonperiodic condition (*M* = .038 μV, *SD* = .084 μV), *t*(17) = - 9.64, p < .0001, *d* = 2.27. This same pattern held when we inspected the category-selective response as a function of ROI. All three regions showed a strong category-selective response in the periodic condition (all 13 harmonics of 0.67 Hz considered significant at p < .001, one tailed), but no such response in the nonperiodic condition (shown in Figure 2 for lateral ROIs).

**Figure 2.**
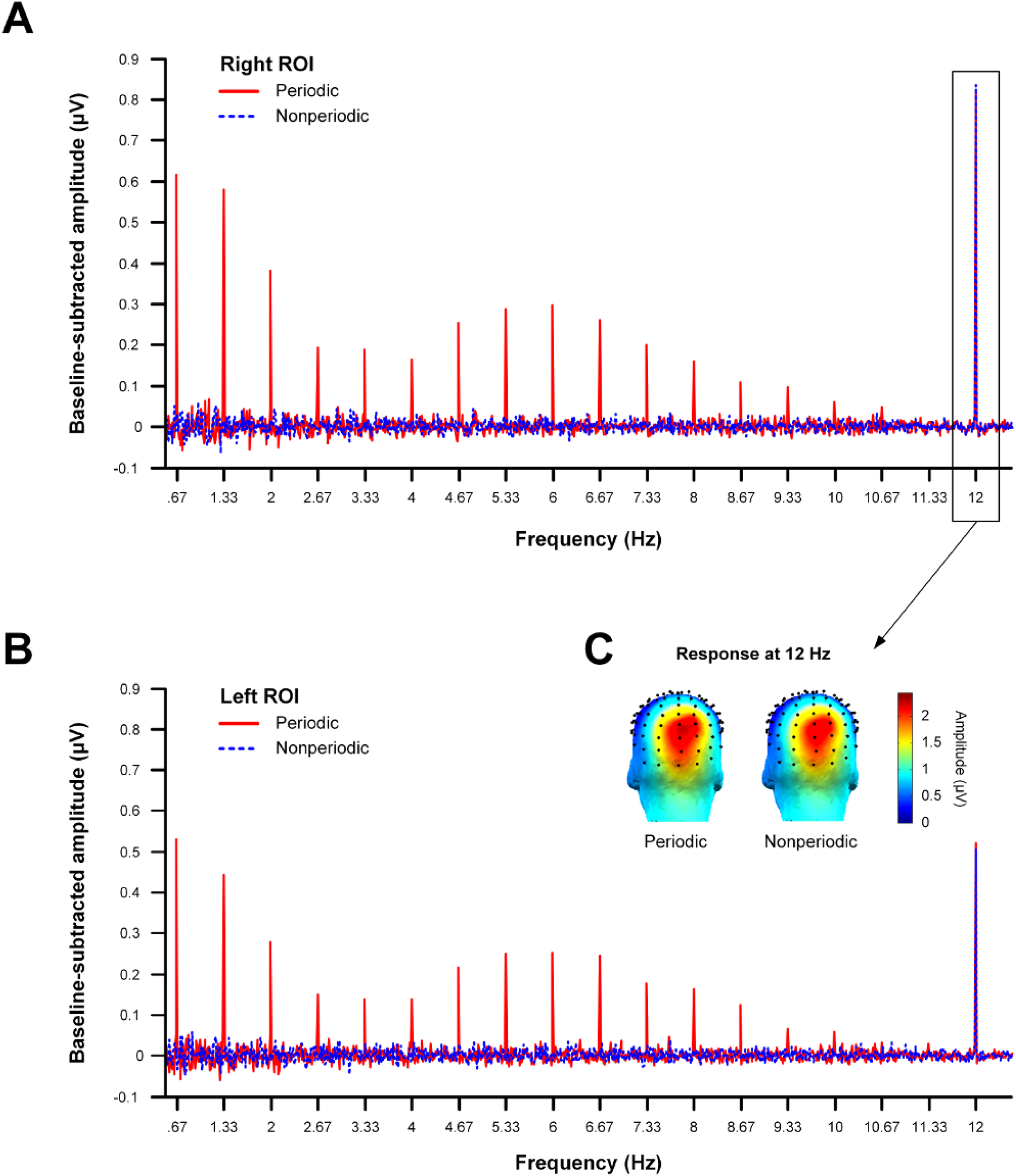
Baseline-corrected amplitude spectra for Expt. 1, shown here as a function of face periodicity for the **(A)** right occipito-temporal region-of-interest (ROI) and **(B)** left ROI. In both regions, the response in the periodic condition at the first 13 harmonics of the category-selective frequency (0.67 Hz) met our criterion for significance (*p* **<** .001, one-tailed). In contrast, there was no response at any of these harmonics in the nonperiodic condition for either ROI. Importantly, both conditions yielded a significant response at the common frequency (i.e., 12 Hz) in both lateral ROIs. **(C)** The response at the common frequency was predominantly centered over medial occipital sites (around Oz).

We quantified the category-selective response in each ROI/condition pair by summing the baseline-corrected amplitude values across the first 13 harmonics of 0.67 Hz (Retter and Rossion 2016), and subjecting these values to a two-way within subjects ANOVA with the factors Periodicity and ROI. Here we observed a significant interaction, *F*(2,34) = 23.86, *p* < .0001, the nature of which is clear in Figure 3. Where ROI clearly modulated the category-selective response in the periodic condition, it had no such impact in the nonperiodic condition, since the responses in this condition were at floor level. Follow-up t-tests showed that all three ROIs showed a strong effect of Periodicity (*p* < .0001 in all cases), with higher responses in the periodic compared to nonperiodic conditions. Owing to the presence of a significant interaction, we did not interpret main effects in this analysis.

**Figure 3.**
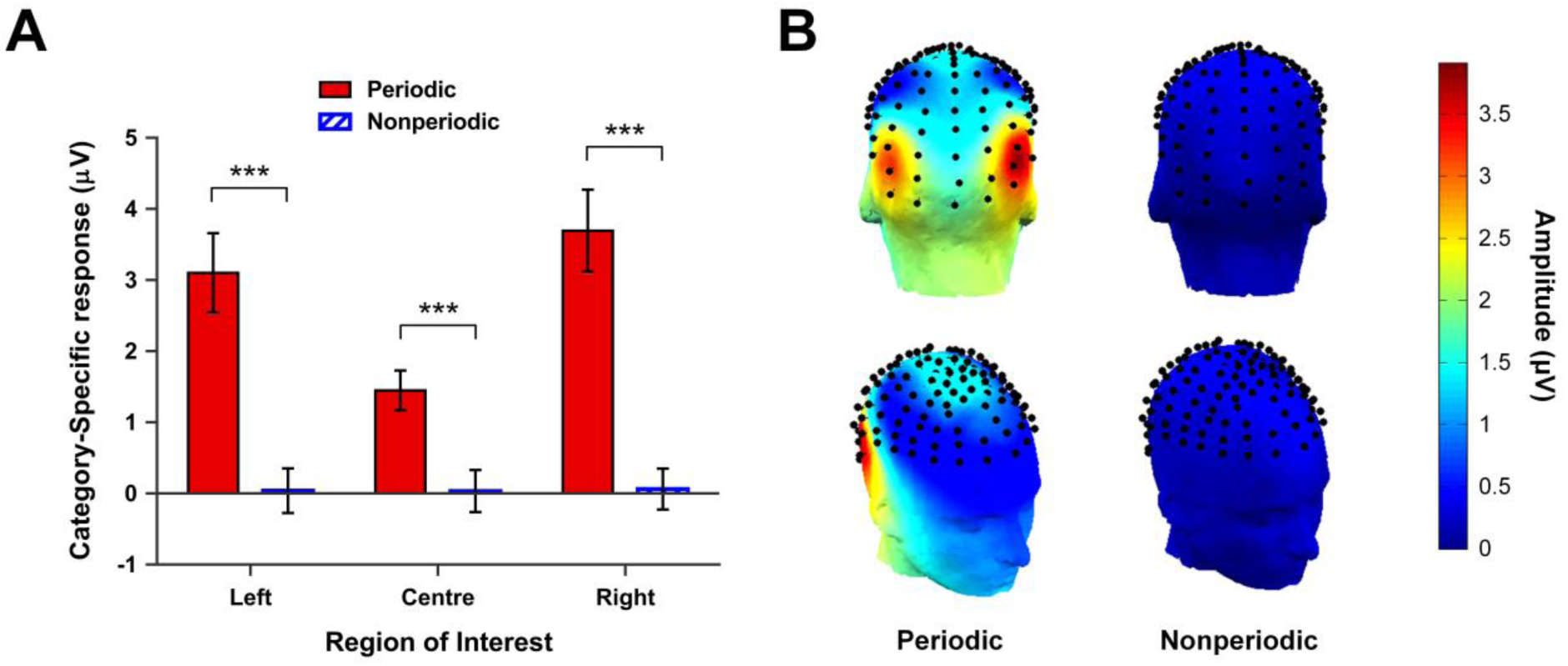
The category-selective response in Expt. 1. **(A)** Sum of baseline-corrected amplitudes at the first 13 harmonics of 0.67 Hz, shown as a function of face periodicity and ROI. All three ROIs showed a strong category-selective response in the periodic condition, but not the nonperiodic condition. Error bars are within-subjects 95% CIs (overlap should not be interpreted by eye, see Cumming & Finch, 2005). Significance codes: ^***^ *p* < .001 **(B)** Scalp topographies for the category-selective response in the periodic and nonperiodic conditions. Where the periodic condition was characterized by a strong bilateral occipito-temporal category-selective response, there was no significant category-selective response at any electrode site in the nonperiodic condition.

#### 3.2.2. Response at the common frequency (12 Hz)

Collapsing across all conditions and channels, we observed a large response at the exact frequency of stimulation (i.e., 12 Hz) and its harmonics 24 Hz and 36 Hz. The scalp response at these frequencies peaked over medial occipital sites (around Oz, see Figure 2C), a topography that is similar to previous studies at this stimulation rate (e.g., Retter and Rossion 2016). Since this response reflects neural synchronization to the stimulus onset/offset rate, and is common to the face and object stimuli, it should not differ between the periodic and nonperiodic conditions here. Indeed, the global scalp response at these three harmonics of 12 Hz was significant for both the periodic and nonperiodic conditions *(p* < .001, one tailed). Moreover, the sum of baseline-corrected amplitude values at these three frequencies did not differ between the periodic (M = 0.713 µV, *SD* = 0.35 µV) and nonperiodic conditions (M = 0.685 µV, *SD* = 0.28 µV), *t*(17) = .519, *p* = .610, *d* = 0.122. Taken together, these results provide no evidence that participants attended differently to the periodic or nonperiodic FPVS sequences.

When we inspected the common response as a function of ROI, we found that all three ROIs showed a highly significant response *(p* < .001, one tailed) at the first three harmonics of the common frequency (i.e., 12, 24, and 36 Hz). Accordingly, we summed across these frequencies and subjected these values to a two-way within subjects ANOVA with the factors *Periodicity* and *ROI.* Here there was no significant interaction, *F*(2,34) = 0.41, *p* = .667, and no main effect of *Periodicity, F*(1,17) = 0.168, *p* = .687. This suggests that the magnitude of the common response was comparable across the periodic (*M* = 1.33 μV, *SD* = 1.04 μV) and nonperiodic (*M* = 1.30 μV, *SD* = 0.93 μV) conditions. In contrast, there was a highly significant effect of *ROI, F*(2,34) = 36.39, *p* < .0001. Follow-up Tukey contrasts showed that the common response was significantly larger in the central ROI (*M* = 2.25 μV, *SD* = 1.08 μV) compared to both the left ROI (*M* = 0.66 μV, *SD* = 0.31 μV), *z* = −8.42, p < .001, *d* = 1.720, and right ROI (*M* = 1.04 μV, *SD* = 0.52 μV), z = −6.37, *p* < .001, *d* = 1.23. In contrast, the common response in the two lateral ROIs did not differ significantly, *z* = 2.04, *p* = .102, *d* = 0.795.

### 3.3. False-Sequencing

Having verified that the periodicity manipulation was effective, we next turned to *quantifying* the response elicited by periodic and nonperiodic faces using a false-sequencing approach (see Methods for details). As for the initial frequency-domain analysis, here too we inspected the category-selective response (imposed in these false-sequences at exactly 1 Hz and its harmonics) as well as the common response at 12 Hz and harmonics.

#### 3.3.1. False-Sequence response at the category-selective frequency (1 Hz).

Using scalp-averaged false-sequence data, we extracted z-scores for the imposed face frequency of 1 Hz and its harmonics under 12 Hz. In both the periodic and nonperiodic false-sequences, all harmonics in all ROIs were highly significant (*p* < .001, one tailed). To compare the magnitude of this whole-scalp category-selective response across-conditions, we summed the response across the first 11 harmonics of 1 Hz. A paired t-test showed there was no significant difference in response magnitude between the periodic (*M* = 1.156 μV, *SD* = .515 μV) and nonperiodic (*M* = 1.104 μV, *SD* = .542 μV) false-sequence conditions, *t*(17) = .836, *p* = .415, *d* = 0.197. Next, we extracted the baseline-subtracted amplitude spectra for the periodic and nonperiodic false-sequences for each of our ROIs. Importantly, all three regions showed a strong category-selective response for both types of false-sequence. That is, in each ROI, all 11 harmonics of 1 Hz were significant at *p* < .001 (one tailed) irrespective of whether the false-sequenced data came from the periodic or nonperiodic condition (see Figure 4).

**Figure 4.**
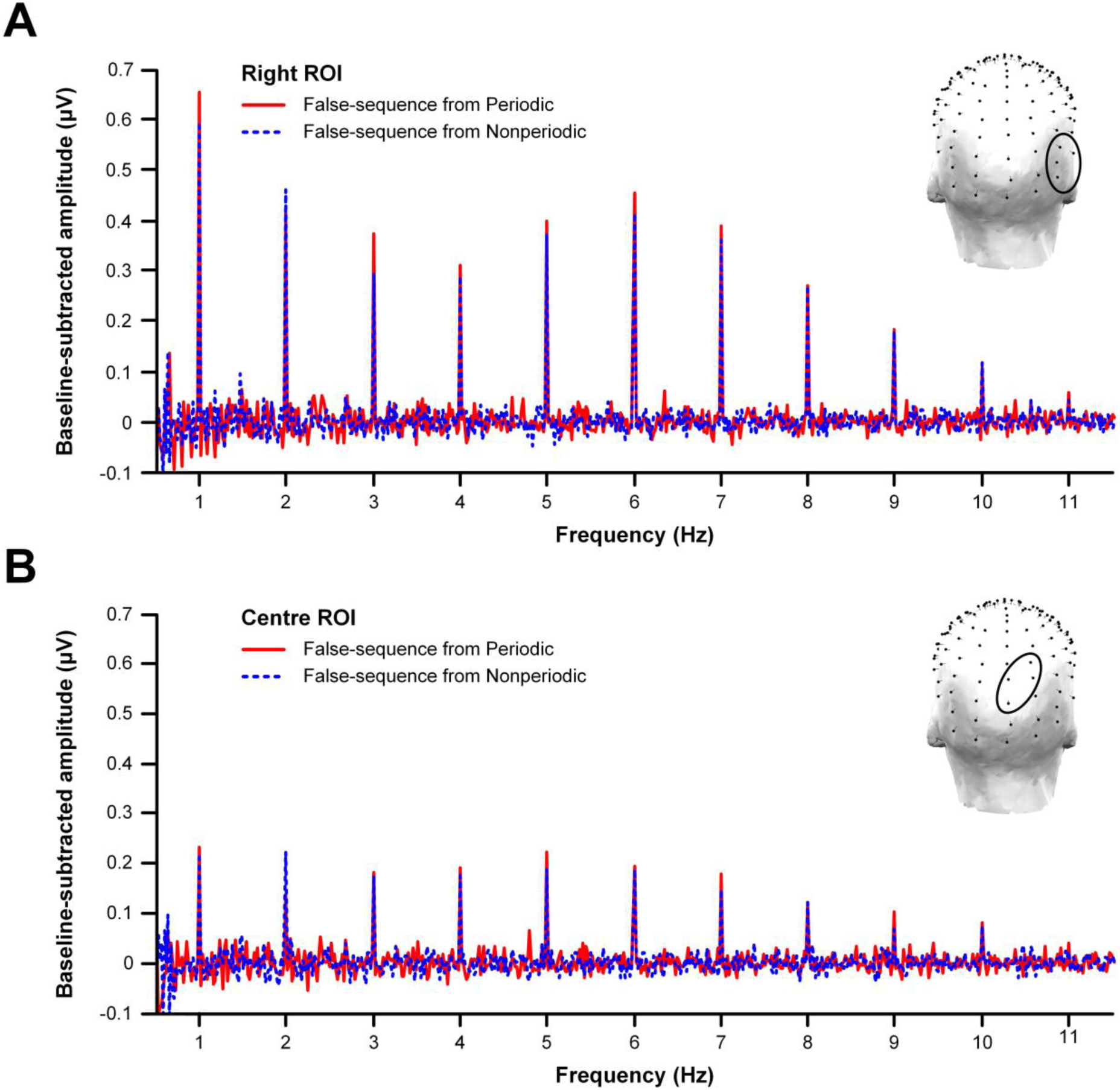
Baseline-subtracted amplitude spectra for the false-sequences corresponding to the periodic and nonperiodic conditions, shown here for the **(A)** right ROI and **(B)** center ROI. Note that unlike the original frequency-domain analyses, the false-sequences generated from both periodic and nonperiodic trials showed a significant response (*p* < .001, one tailed) at the category-selective frequency (i.e., 1 Hz) and its harmonics.

To quantify the response to periodic and nonperiodic faces, we summed the baseline-corrected amplitude values at the 11 harmonics for each condition/ROI pair (see Figure 5A), and subjected these data to a two-way within-subjects ANOVA with the factors *Periodicity* and *ROI*. The interaction between these factors did not approach significance, *F*(2,34) = 1, *p* = .379, nor did the main effect of *Periodicity*, *F*(1,17) = 2.79, *p* = .113. As such, there was no evidence that category-selective response differed between the periodic (*M* = 2.76 μV; *SD* = 1.55 μV) and nonperiodic false-sequences (*M* = 2.58 μV; *SD* = 1.61 μV). In contrast, there was a significant main effect of *ROI, F*(2,34) = 22.87, *p* < .0001, which we followed up with Tukey contrasts. As can be seen in Figure 5A, the response in the right ROI (*M* = 3.51 μV, *SD* = 1.57 μV) was greater than the response in both the center ROI (M = 1.65 μV; *SD* = .835 µV), *z* = 6.86, *p* < .001, *d* = 1.508, and left ROI (*M* = 2.85 µV; *SD* = 1.61 μV), z = 2.43, *p* = .04, *d* = 0.612. Also, the response in the left ROI was greater than that in the center ROI, z = 4.43, *p* < .001, *d* = 0.975.

**Figure 5.**
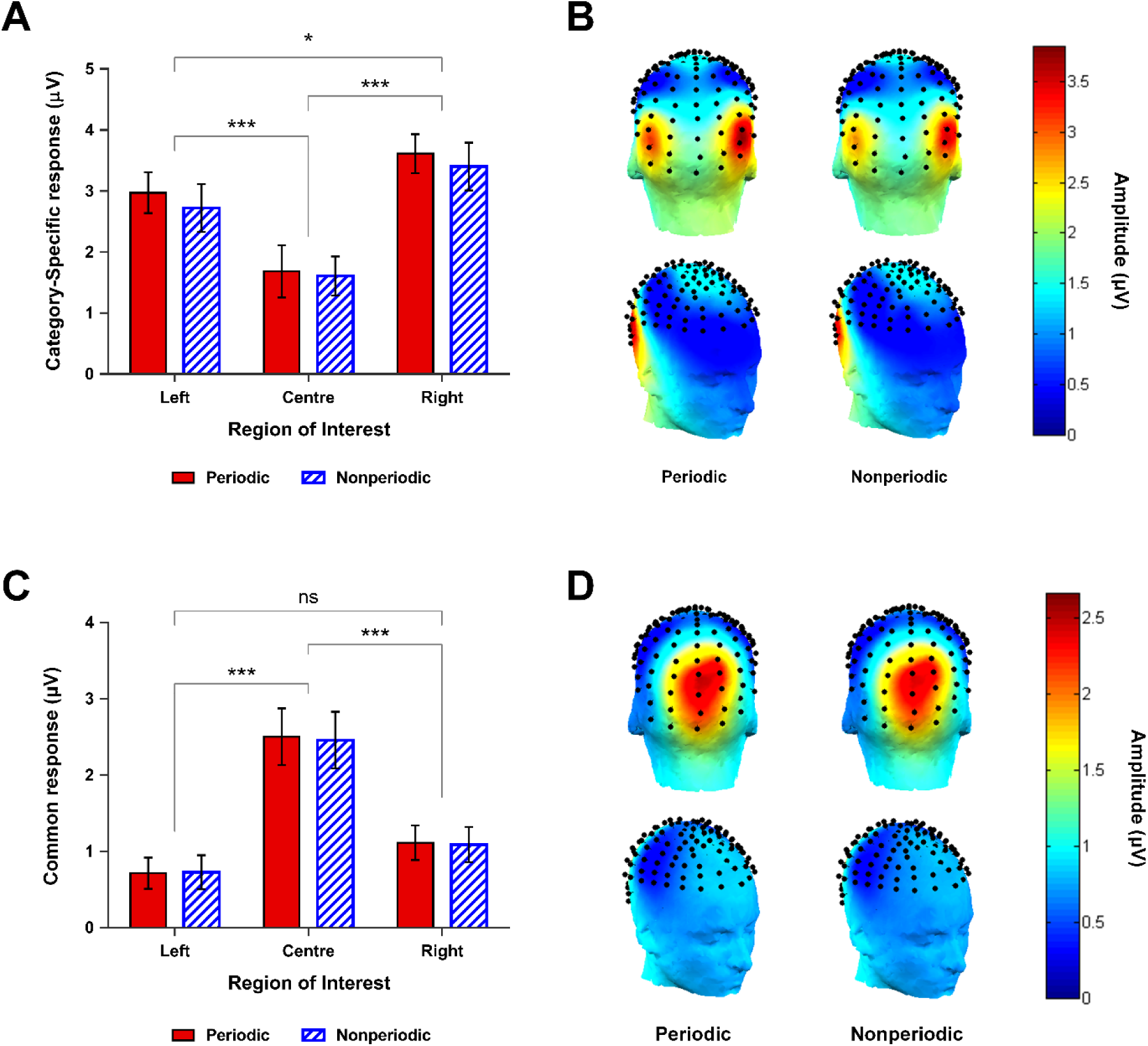
Sum of baseline-corrected amplitudes representing the **(A)** category-selective response (first 11 harmonics of 1 Hz) and **(C)** common response (first 3 harmonics of 12 Hz) for the false-sequence data in Expt. 1. Both the category-selective and common response were strongly modulated by ROI, but were not at all sensitive to face periodicity. Error bars are within-subjects 95% CIs (overlap should not be interpreted by eye, cf. Cumming, 2005). Significance codes: ^***^ ***p*** < .001; ^**^ ***p*** < .01; ^*^ ***p*** < .05. **(B)** Scalp topographies corresponding to the category-selective response and **(D)** common response, shown as a function of face-periodicity.

Since we were conscious of the problems associated with drawing inferences from *p*-values in support of the null hypothesis (Dienes 2014), we also calculated the Bayes factor for each model combination involving the factors Periodicity and ROI. This analysis showed that the ROI-only model provided the best fit, accounting for our data approximately 29,620 times better than the null model (intercept only). In contrast, the Bayes factor for the *Periodicity*-only model was just 0.238, suggesting the null model accounted substantially better for our data than one including just *Periodicity* (Jeffreys H 1939/1961). Furthermore, when comparing our preferred *ROI*-only model with all other model combinations, it performed approximately 4.05 times better than the next best model which included additive effects of both *ROI* and *Periodicity.* Thus, the inclusion of *Periodicity* did not add sufficient explanatory power to overcome Bayesian penalties for increasing model complexity.

Given the similarity in the magnitude of the response to periodic and nonperiodic faces, we expected there should be a strong correlation between the sum of category-selectiv harmonics for the periodic and nonperiodic condition across participants. Figure 6 shows clearly that this was indeed the case - there was a very high correlation between the face response for the periodic and nonperiodic conditions in both the left ROI, *r*(16) = .931, *p* < .0001, and the right ROI, *r*(16) = .935, *p* < .0001. In short, individuals with a strong face-selective response in the periodic condition also had a strong face-selective response in the nonperiodic condition.

**Figure 6.**
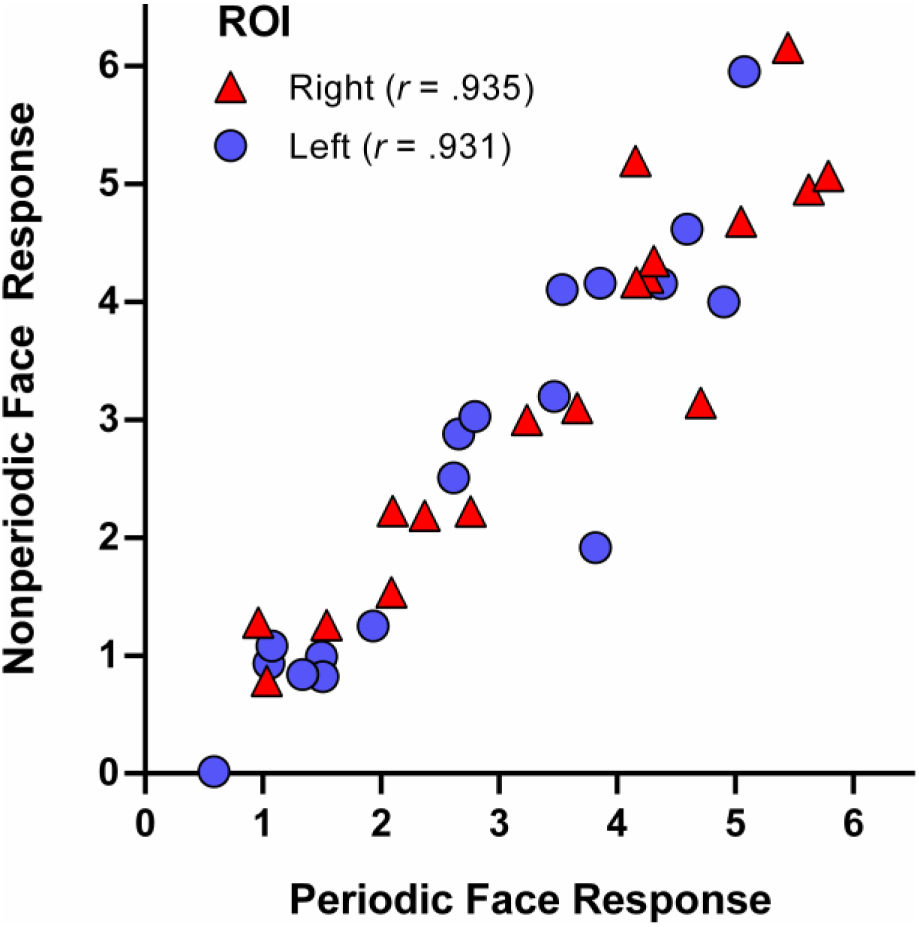
The correlation between the category-selective response (summed baseline-corrected amplitudes across first 11 harmonics of 1 Hz) for the periodic and nonperiodic false-sequences. In both the right ROI (red triangles) and left ROI (blue circles), we observed a strong correlation in category-selective response magnitude between the periodic and nonperiodic false-sequences.

#### 3.3.2. False-Sequence response at the common frequency (12 Hz)

Where the false-sequencing procedure imposes the category-selective response at a defined frequency (here at 1 Hz), it should have no impact on the frequency at which the common response is observed (i.e., 12 Hz). Indeed, collapsing across all channels and conditions, our z-score extraction procedure identified a highly significant response for both false-sequence types at 12 Hz, 24 Hz, and 36 Hz (*p* < .001, one tailed). Accordingly, we subjected the sum of baseline-corrected amplitudes at these three frequencies for each condition/ROI pair to a within-subjects two-way ANOVA with the factors *Periodicity* and *ROI*. Here we observed an identical pattern of results to that seen in the original frequency-domain analysis (see Figure 5B). That is, the *Periodicity* x *ROI* interaction was not significant, *F*(2,34) = 1.83, *p* = .176, nor was the main effect of *Periodicity, F*(1,17) = .877, *p* = .362. Thus, there was no evidence that the magnitude of the common response differed for periodic (*M* = 1.44 μV; *SD* = 1.10 μV) and nonperiodic (*M* = 1.42 µV; *SD* = 1.09 μV) false-sequences. In contrast, there was a strong main effect of *ROI, F*(2,34) = 32.63, *p* < .0001. Follow-up Tukey contrasts indicated this was due to significantly lower responses in the left *ROI* (*M* = .720 μV; *SD* = .303 μV) and right *ROI (M* = 1.10 μV; *SD* = .519 μV) compared to the center ROI (*M* = 2.48 μV; *SD* = 1.23 μV), Center-Left: *z* = −7.90, *p* < .0001, *d* = 1.565; Center-Right: z = −6.19, *p* < .0001, *d* = 1.191. In contrast, the common response in the right and left ROIs was not significantly different, z = 1.71, *p* = .203, *d* = 0.790.

### 3.4. Time-Domain Results

We further examined the response elicited by periodic and nonperiodic faces in the time-domain. Figure 7 shows the mean waveforms for all 128 channels time-locked to the onset of periodic and nonperiodic faces. This response was remarkably similar in the two periodicity conditions, notably characterized by the same four distinct spatiotemporal components we have reported previously for a different set of faces in an FPVS-EEG design (Retter and Rossion 2016).

**Figure 7.**
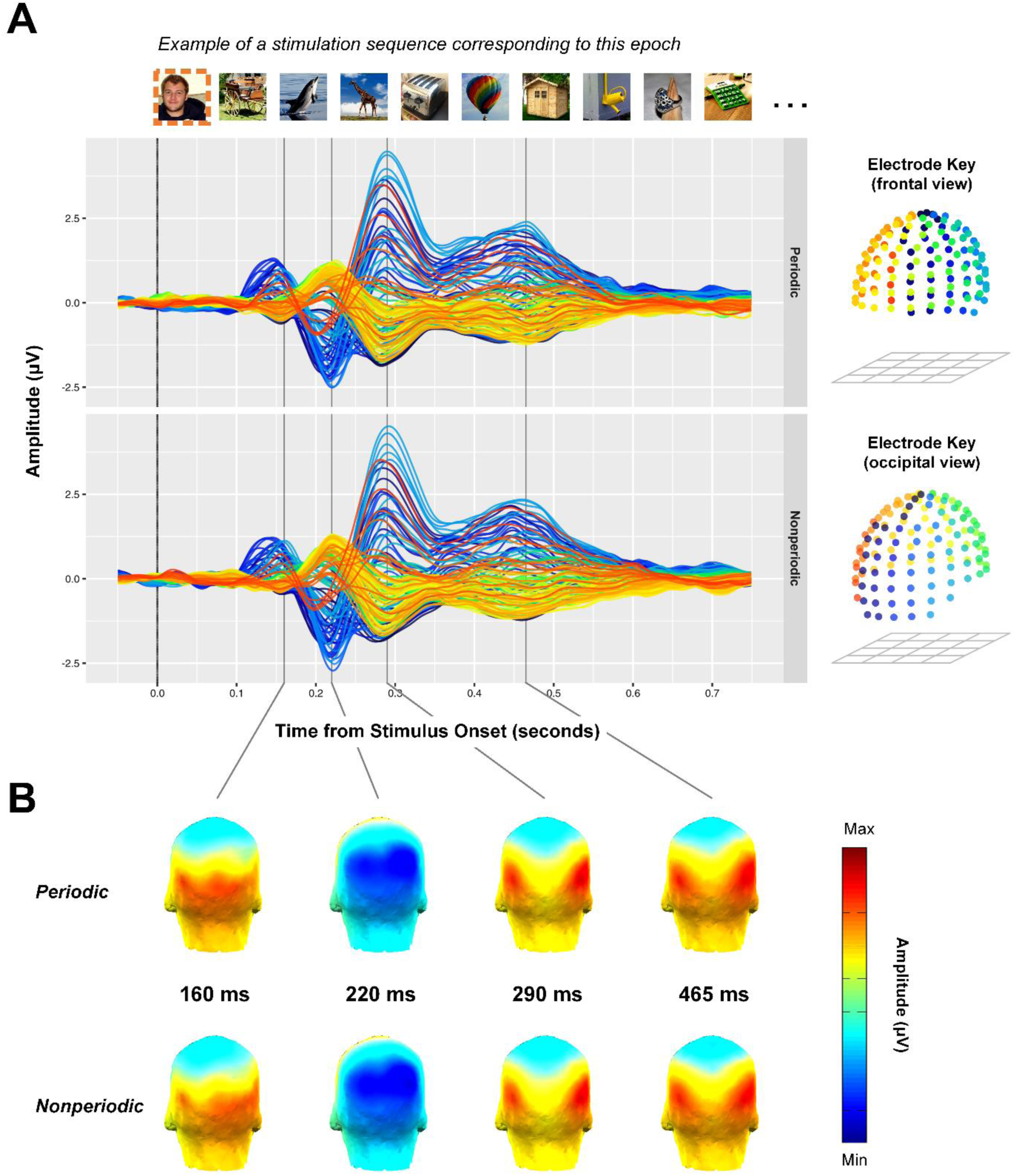
Time-domain representation of the category-selective response elicited by faces in the periodic and nonperiodic conditions. **(A)** Conditional mean waveforms for all 128 channels plotted as a function of time from face onset. The image presentation frequency (i.e., 12 Hz) has been removed here using notch-filtering, see Figure 12, also Retter & Rossion, 2016, Figure 4). **(B)** Scalp topographies for the periodic and nonperiodic conditions corresponding to the four spatiotemporal peaks comprising the category-selective response (0.165 s, 0.220 s, 0.290 s, and 0.465 s). Amplitude scales are fixed for each condition pair at each time points.

We compared the waveforms elicited by temporally predictable and unpredictable faces in each ROI by conducting a paired *t*-test at each of the 513 samples. Any observed *t*- value that fell outside the critical cut-off given by our conservative permutation procedure (see Methods) was considered a significant difference between the periodic and nonperiodic conditions. As can be seen in Figure 8, no observed *t*-value in any ROI approached either this permutation defined cut-off, or even a less conservative cut-off of *p* < .01 (uncorrected). As such, there was no evidence to suggest that the response evoked by periodic and nonperiodic faces in the fast periodic visual stream differed meaningfully in any ROI.

**Figure 8.**
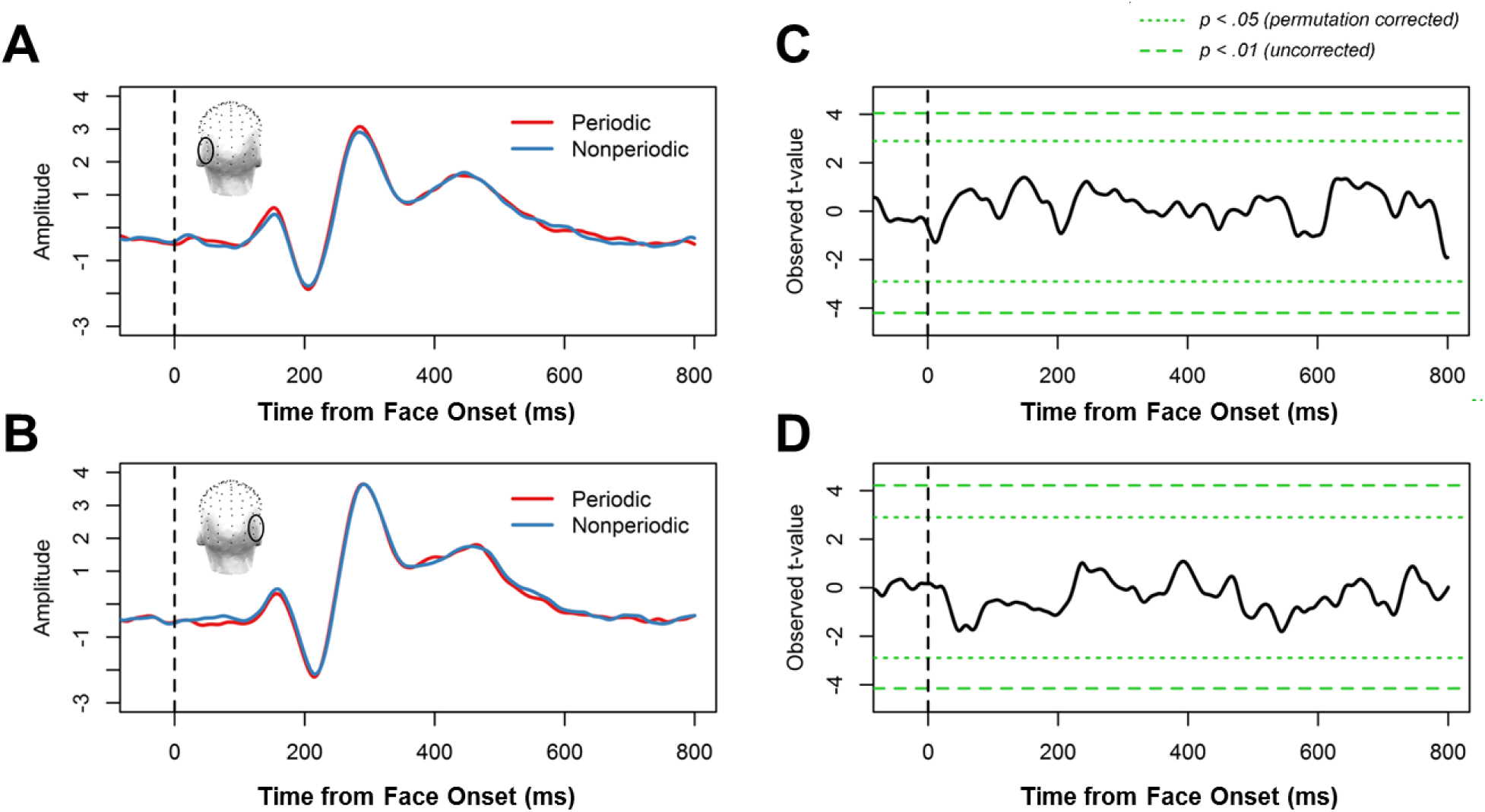
Time-domain analysis results for Expt. 1. Left column: Conditional mean amplitude following 12 Hz notching-filtering as a function of time from face onset (vertical dashed line), shown for the **(A)** left ROI and **(B)** right ROI. Right column: The observed *t*-value for each time point, shown for the **(C)** left ROI and **(D)** right ROI. Horizontal dashed lines reflect criterion significance cut-offs (permutation corrected and uncorrected). Note that no observed *t*-value at any sample in any ROI approached either criterion cut-off value, suggesting there was no difference in the response evoked by periodic and nonperiodic faces.

To further assess any difference in the response evoked by periodic and nonperiodic faces, we subjected our time-domain data to MVPA decoding. In the first of these analyses we trained an SVM classifier on the spatiotemporal activity patterns associated with periodic faces and nonperiodic faces in 20ms increments (-166ms to 500ms, see Methods for details). Paired *t*-tests (*n* = 34) indicated that this classifier was not able to identify novel, unlabeled face responses as either periodic or nonperiodic significantly better than empirical chance at any time point following face onset. A complementary question was whether a classifier trained on periodic face responses would generalize well to nonperiodic face responses. To this end, we trained a second SVM classifier on spatiotemporal patterns of activation elicited by face and object images in the periodic condition (-166ms to 500ms, see Methods for details). We then tested classifier performance using novel data drawn from either the periodic condition (i.e., within-condition decoding), or the nonperiodic condition (i.e., cross-condition decoding). Here classifier decoding accuracy for the within-condition analysis acts as a baseline, reflecting the degree to which the response to faces can be reliably distinguished from the response to objects. Indeed, within-condition decoding of face and object responses was significantly higher than empirical chance from ~137 ms to ~352 ms following stimulus onset, with peak decoding at ~274 ms post onset (Figure 9A, blue lines). Importantly, cross-condition decoding accuracy was also significantly above empirical chance from ~118 ms to ~371 ms after stimulus onset, with peak decoding at exactly the same point (~ 274 ms) as in the within-condition decoding analysis (Figure 9B, red lines). Moreover, paired *t*-tests indicated decoding accuracy was not significantly different in the within-condition and cross-condition analyses at any time, suggesting the response evoked by faces in the periodic condition generalized equally well to nonperiodic novel data as to periodic novel data. Indeed if anything, cross-condition decoding started slightly earlier and lasted longer than within condition decoding, even though there was no significant difference in decoding accuracy at any time point.

**Figure 9.**
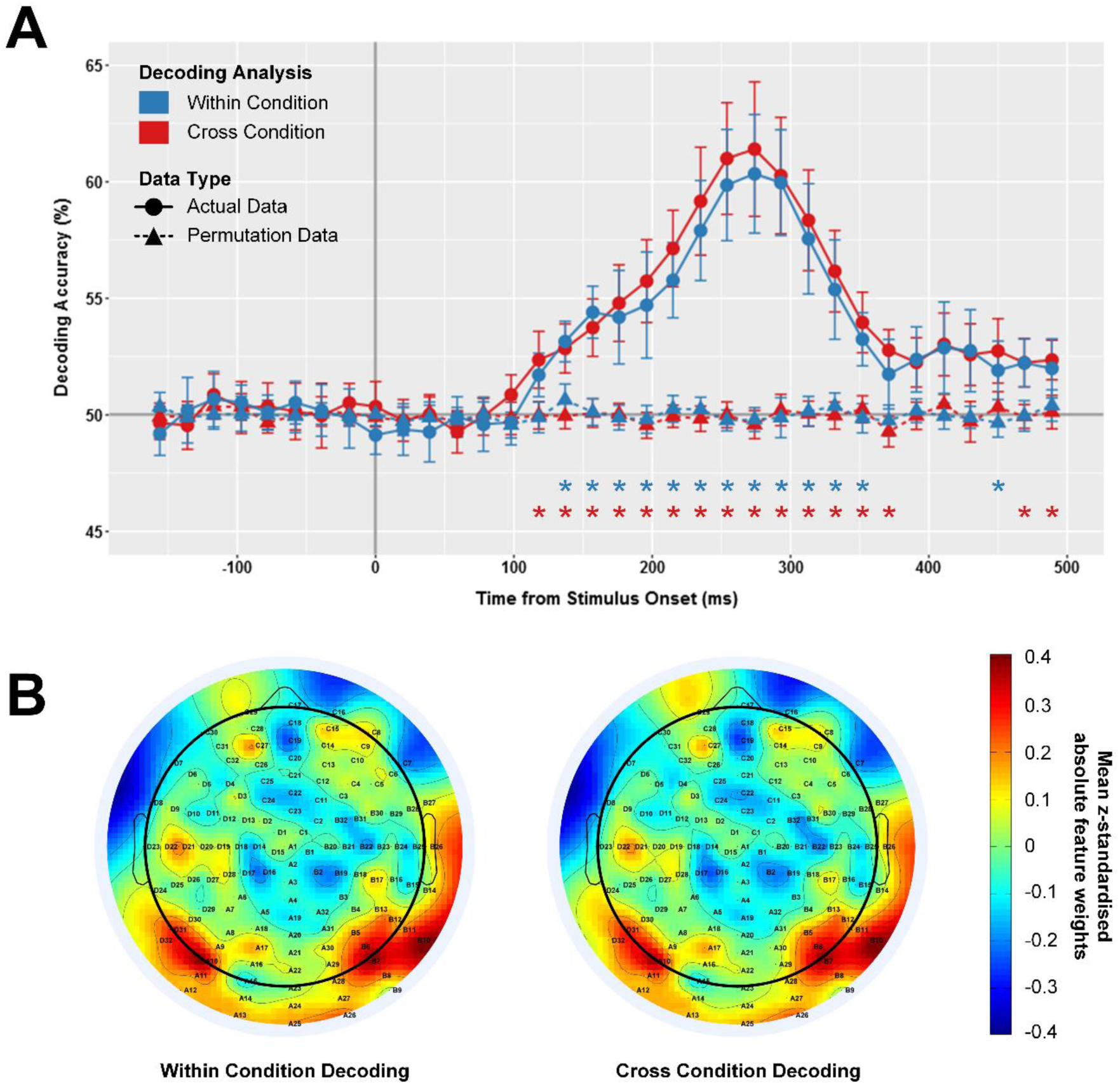
**(A)** Face vs. object decoding accuracies for the within-condition (blue) and the cross-condition (red) analyses, shown as a function of data type (actual and permutated). Error bars represent SEM; asterisks indicate significantly higher decoding accuracy for the actual data compared to the permutated control data. Note that decoding accuracy was not significantly different for the within-condition and cross-condition analyses at any time point **(B)** z-standardized absolute feature weights for all 128 channels, averaged across 150ms to 350ms, shown for the within-condition and cross-condition analyses. In both cases, the primary sources of face vs. object information were bilateral occipito-temporal channels.

## 4. Experiment 1 Interim Discussion

In Expt.1 we set out to compare the category-selective response evoked by faces embedded at periodic or nonperiodic intervals in an FPVS sequence. We found no evidence of a qualitative or quantitative difference in the response to temporally predictable and unpredictable faces. In both conditions, the category-selective response for faces was comprised of the same four spatiotemporal components we have reported previously for another set of natural face images in FPVS sequences (Jacques et al. 2016; Retter and Rossion 2016): P1-face (peaking at 145 ms), N1-face (peaking at 220 ms), P2-face (peaking at 290 ms), and P3-face (peaking at 465 ms). Several results point to the conclusion that the category-selective response, of about 400ms total duration (Retter & Rossion, 2016), was immune to the temporal predictability of faces in the rapid sequence. First, our quantification analysis of the response to periodic and nonperiodic faces (false-sequencing approach) showed that the magnitude of the face-selective response was not sensitive to face periodicity, although it was modulated by ROI (right > left > center). Moreover, Bayesian statistics indicated that a null model was actually preferred over one containing face periodicity as a factor. Second, permutation test procedures across the 800 ms following face onset found no difference in the response to periodic and nonperiodic faces in any of our ROIs, nor was a linear classifier trained on periodic and nonperiodic face responses able to classify unlabeled responses better than empirical chance. Importantly, however, a different classifier trained to differentiate the response evoked by faces and objects in the periodic condition was able to classify unlabeled face/object responses equally well regardless of whether they were drawn from the periodic or nonperiodic condition. This latter result suggests that the face-selective response was qualitatively similar in the two periodicity conditions. Taken together, these results strongly suggest that temporal predictability of critical category exemplars in FPVS sequences does not generate, or even modulate, the category-selective response in these designs.

Why did periodic and nonperiodic faces in our study elicit such similar responses, given that several existing studies have reported that rhythmic temporal stimulation does modulate perceptual processing (Cravo et al. 2013; Breska and Deouell 2014; Morillon et al. 2016)? One possibility is that an effect of temporal expectation was masked here by an overall attentional imbalance between the conditions. Specifically, if observers paid more attention to the FPVS sequences in one condition compared to the other, then the category-selective signal in the better-attended condition could have been boosted so that it equaled that in the other condition, thereby concealing an effect of temporal predictability. If that were the case, however, there should be a commensurate boost to the response at the common frequency as well (i.e., 12 Hz), given that such “steady-state visual evoked potential” (SSVEP) responses at these high frequency rates are substantially increased by selective attention over medial occipital sites (e.g., Müller et al. 2006). We found no such difference in the magnitude of the 12 Hz response between conditions, suggesting participants attended to the sequences equally well regardless of face periodicity. As such, a difference in attentional allocation seems unlikely to account for why temporal predictability did not modulate the category-selective response in our study.

An alternative possibility is that, owing to the high saliency of face stimuli for the human brain (Hershler and Hochstein 2005; Crouzet et al. 2010), participants’ temporal expectations about the predictable onset of faces were not sufficient to modulate the strong face-selective evoked response. That the neural response to faces might be robust to modulation by temporal expectation is not an unreasonable possibility, considering that effects of factors such as spatial attention on face-processing have historically been quite difficult to find (Reddy et al. 2004; Reddy et al. 2006; Finkbeiner and Palermo 2009; Quek and Finkbeiner 2013). If is indeed the case, then another way to examine whether temporal predictability drives the category-selective response in FPVS is to observe the neural response to a violation of temporal expectations, i.e., when the visual stimulus encountered does not match a pre-activated face template (Rao and Ballard 1999; Friston 2005; Kok et al. 2014). We tested this hypothesis in Experiment 2.

## 5. Experiment 2

To examine the neural response to a violation of rhythmic temporal expectations, participants in Expt. 2 viewed rapid continuous sequences of natural object images with a face embedded as every 12^th^ image (i.e., at periodic intervals in the sequence). After a period of entrainment, a small fraction (10 %) of these highly temporally predictable faces were replaced with a randomly selected object image, a stimulus we refer to here as a “missing face”. If temporal expectations about the critical stimulus category contribute to the category-selective signal in FPVS, then non-face stimuli that violate these expectations (i.e., missing faces) should elicit a different response than non-face stimuli about which participants cannot build any temporal expectations. In simple terms, the response evoked by an object image that appears in place of an expected face should differ in some way from that evoked by the other object images in the sequence. If this is indeed the case, a false-sequencing analysis of missing faces should capture this differential response. Importantly, the false-sequencing approach makes no assumptions about what the nature of the differential response to missing faces should be. It could be, for example, that objects replacing expected faces evoke a “prediction error” type response, perhaps similar in kind to the mismatch negativity (MMN; for a review, see Naatanen et al. 2010) that has been associated in recent theoretical accounts with an error detection signal (Friston 2005; Garrido et al. 2009; Kimura et al. 2011; Stefanics et al. 2011; Lieder et al. 2013; Pieszek et al. 2013). On the other hand, perhaps participants’ temporal expectations about faces actually drive a partly face-like selective response to objects that replace expected faces, even in the absence of a true face stimulus. Previous studies have shown, for example, that unexpected omissions of visual stimuli can elicit feature-specific responses in visual cortex (den Ouden et al. 2009; Kok et al. 2014). Whatever its exact nature, if the response evoked by objects replacing expected faces differs in any way from that evoked by other objects in the sequence, it will be reflected in a significant signal at the imposed frequency in a false-sequence.

## 6. Experiment 2 Methods

### 6.1. Participants

A different group of 13 individuals (eight females) aged between 19-26 years took part in Expt. 2 in exchange for payment. Participation criteria were the same as Expt. 1; we obtained written informed consent from all participants prior to testing.

### 6.2. Procedure

The stimuli, experimental testing set up, and fixation cross task were identical to those described in Expt. 1. Each trial consisted of a pre-stimulation period which showed only the central fixation cross (2 s), a fade-in period (2 s), the stimulation sequence proper (120 s), and a fade-out period (3 s). As in Expt. 1, the stimulation sequence was a series of object images shown at a periodic rate of 12 Hz. Here we embedded the sequence with a natural face image after every 11 object images, giving a face periodicity of exactly 1 Hz (i.e., 12 Hz image presentation rate / 12 images). The critical manipulation was the inclusion of “missing faces” within each sequence, achieved by replacing 10% of the periodic faces with a randomly selected object image (see Figure 10)^1^. Replacements could only occur after an initial entrainment time of at least 10 s (i.e., 10 normal face presentation cycles). We separated the missing faces by a variable delay of between 6 to 15 normal face presentation cycles (i.e., 6-15 s). Thus, each sequence contained approximately 108 real faces and up to 12 missing face instances, so that the conditions were optimal for temporal expectation. Participants viewed 10 sequences in total.

**Figure 10.**
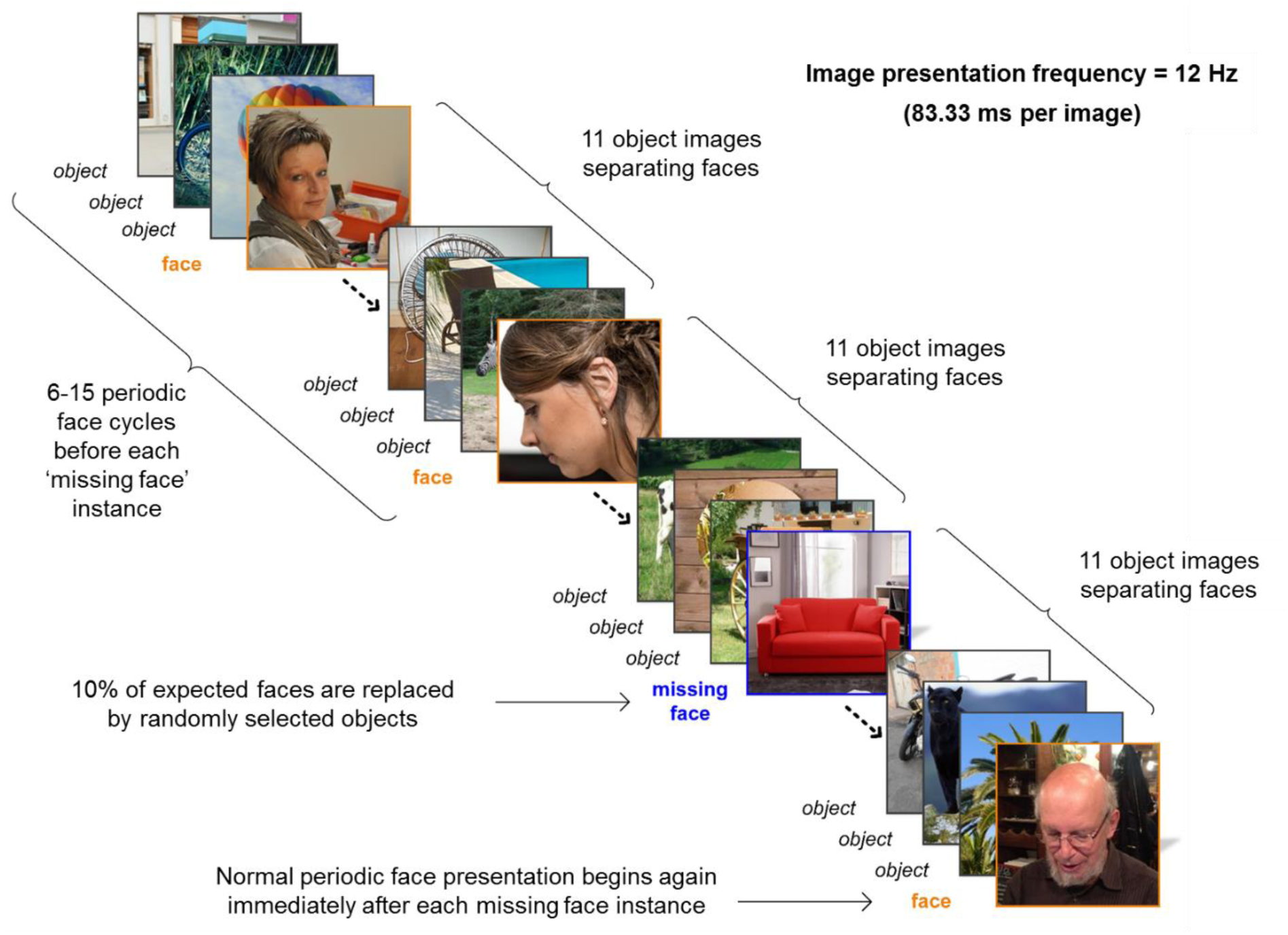
Schematic representation of the stimulation sequence for Expt. 2. Natural object images appeared at a periodic rate of exactly 12 Hz (i.e., 12 images per second), with a face image embedded as every 12^th^ image (face periodicity = 1 Hz). On each trial, 10% of these periodic faces were replaced by a randomly selected object image, resulting in a “missing face” for the observer. Between six to 15 normal periodic face cycles occurred between such missing face instances. Throughout the sequence, the participant’s task was to monitor a central fixation cross which overlaid the images for changes of color.

### 6.3. EEG Analysis

#### 6.3.1. EEG pre-processing

We excluded data from two female participants whose EEG trace on initial inspection contained excessive artefact that unduly affected time-domain visualization. As in the case of the participants removed in Expt. 1, both individuals nevertheless showed a significant response at the face periodicity and several harmonics (1 Hz). All remaining pre-processing steps were identical to those implemented in Expt. 1, save that here we segmented longer epochs to correspond to the 120 s stimulation sequences.

#### 6.3.2. Frequency-domain analysis

The goal of the initial frequency analysis was to verify that sequences in which 10% of the periodic faces were removed nevertheless still evoked a category-selective response (i.e., response at 1Hz) indicating participants synchronized to the periodic rate of face stimulation. To this end, we cropped the EEG recording for each stimulation sequence to a 119.02 second epoch containing exactly 119 face presentation cycles, and averaged the 10 epochs for each participant before applying an FFT. The frequency resolution of the resulting normalized amplitude spectra was 0.0084 Hz (the inverse of the sequence duration, i.e., 1/119.02). As in Expt. 1, we identified significant responses at the frequencies of interest by pooling the amplitude spectra across channels and computing a *z*-score at each frequency bin (significance criterion = *z* > 3.1, i.e, *p* < .001, one-tailed). We then quantified the response (sum of baseline-corrected amplitude) at the relevant frequencies in three ROIs defined using the same procedures as for Expt. 1. This gave left and right occipito-temporal ROIs that were identical to those of Expt. 1 (left: P9, PO7, PO9, & PO11; right: P10, PO8, PO10, & PO12), and a near-identical central ROI (O2, POOz, PO4h, POO6)^2^.

#### 6.3.3. False-sequencing analysis

Our goal here was to quantify and compare the response to real faces and the response to “missing” faces using the same false-sequencing procedure described for Expt. 1. Epochs were 1000ms long, lasting from 500ms before to 500ms after each missing face instance, resulting in ~12 segments that we then sequenced, linearly de-trended, and cropped to exactly 11 s. We repeated this same procedure to produce false-sequences corresponding to each real face occurrence that immediately preceded a missing face. This ensured an equal number of real and missing faces in the false-sequencing analysis, in which the imposed critical stimulus periodicity was exactly 1 Hz. For the two false-sequence types, we de-trended and averaged the epochs before applying an FFT and performing a baseline correction (as in Expt. 1).

#### 6.3.4. Time-domain analysis

We inspected the EEG waveform data time-locked to a comparable number of missing faces and real faces. First, we low-pass filtered the data using a Butterworth with a 30 Hz cutoff and cropped each sequence to be an integer number of 12 Hz cycles. We applied a notch filter (width = P 0.05 Hz) to remove the response at the common frequency of 12 Hz, and three of its harmonics (24 Hz, 36 Hz, and 48 Hz). We then segmented an epoch lasting from −200 ms to 750 ms around *i)* each missing face in the sequence, and *ii)* each real face instance immediately preceding a missing face. We then baseline-corrected each epoch to the average of the 166ms preceding stimulus onset, and averaged the resulting ~120 epochs of each type to create conditional means. A paired *t*-test between the condition of interest and a baseline of zero at each time point (*n* = 564) was computed to assess the presence of a category-selective response for each condition. As in Expt. 1, we inspected the obtained *t*-values relative to two significance criteria (*p* < .01 uncorrected, and *p* <.05, permutation corrected). We obtained the second, more conservative criterion using the same permutation procedure described for Expt. 1 (2,048 permutations implemented across 564 time points, for a total of 1.16 million comparisons).

## 7. Experiment 2 Results

### 7.1. Behavior

Detection accuracy for the fixation cross color change task was near ceiling at 95.85% *(SD* = 6.02%). The mean response time (RT) on correct trials was 502 ms *(SD* = 42ms).

### 7.2. Frequency-Domain Results

#### 7.2.1. Response at the category-selective frequency (1 Hz).

Collapsing across all scalp channels, we found a highly significant (*p* < .001, one tailed, i.e., signal > noise) response at each of the first 18 harmonics of the category-selective frequency (1 Hz). Thus, even though 10% of the periodic faces were missing, the periodic faces nevertheless still evoked a strong category-selective response, indicating that participants did indeed synchronize to the periodic rate of face stimulation. A one-way within-subjects ANOVA indicated that the sum of these 18 harmonics (excluding the 12^th^ harmonic, i.e., the 12 Hz common frequency) varied significantly with *ROI, F*(2,20) = 10.86, *p* < .0001. Follow up Tukey contrasts indicated that the right (*M* = 3.45; *SD* =1.76) and left ROIs (*M* = 2.74; *SD* = 1.71) exhibited a significantly larger category-selective response than the central ROI (*M* = 1.76; *SD* = .904) (right vs. center: *z* = 4.87, *p* < .001, *d* = 1.164; left vs. center: z = 2.82, *p* = .013, *d* = .756). However, these two lateral regions did not differ significantly from one another, *z* = 2.04, *p* = .102, *d* = 0.927.

#### 7.2.2. Response at the common frequency (12 Hz).

Again collapsing across all channels, we also found a highly significant (*p* < .001, one tailed) response at the first five harmonics of the common frequency (i.e., 12 Hz, 24 Hz, 36 Hz, 48 Hz, and 60 Hz). We quantified this response by summing baseline-subtracted amplitude values across these five harmonics in each of our predefined ROIs. A one-way within-subjects ANOVA revealed a highly significant effect of *ROI* on the magnitude of the common response, *F*(2,20) = 25.02, p < .0001. As was the case in Expt. 1, follow up Tukey contrasts showed this response was significantly larger in the central ROI (*M* = 2.13 μV, *SD* = .978 μV) compared to both the left ROI (*M* = .562 μV, *SD* = .246 μV), *z* = −7.09, *p* < .0001, *d* = 1.522, and right ROI (M = .926 µV, *SD* = .426 μV), *z* = −5.44, *p* < .0001, *d* = 1.679. In contrast, there was no significant difference in the magnitude of the common response between the left and right ROIs, *z* = 1.65, *p* =.224, *d* = 0.814.

### 7.3. False-Sequencing

The false-sequencing procedure generated 12 Hz sequences containing either a “missing face” or real face at an imposed periodicity of 1 Hz. We inspected the response at both the category-selective frequency and common frequency for these two sequence types.

#### 7.3.1. False-sequence response at the category-selective frequency (1 Hz).

Pooling across all scalp channels and conditions, we identified significant responses (*p* < .001, one tailed) at the 4-8^th^ harmonic of the imposed periodicity at 1 Hz. As such, we summed the baseline-corrected amplitude values for each condition across the initial eight harmonics of 1 Hz ^3^. Averaged across the whole scalp, this sum of category-selective harmonics was significantly larger for the real face sequences (*M* = .526 μV, *SD* = .391 μV) compared to the missing face sequences (*M* = -.171 μV, *SD* = .150 μV), *t*(10) = 5.60, *p* < .001, *d* = 1.688). We then inspected the sum of harmonics as a function of ROI with a two-way within-subjects ANOVA with the factors Sequence Type and ROI. Here there was a significant interaction between *Sequence Type* and *ROI, F*(2,20) = 4.59, *p* = .023, the nature of which is clear in Figure 11A. As in Expt. 1, where ROI significantly modulated the category-selective response in the real face condition, it did not in the missing face condition (since the response in the latter condition was at floor level). Follow up pairwise t-tests indicated there was a significant effect of Sequence Type in each ROI (all Bonferroni adjusted *p* values < .05). Owing to the presence of a significant interaction between Sequence Type and ROI, we did not inspect the main effects for these factors. However, since we were primarily interested in asking whether the missing faces generated any category-selective response *at all,* we did inspect the 95% confidence interval (CI) around the mean category-selective response for missing faces, and noted that it included zero (-0.493 to .148) where the 95% CI for real faces did not (1.170 to 1.812). This would suggest that real faces gave rise to a category-selective (i.e., differential) response (*M* = 1.49 μV; *SD* = 1.29 μV), where missing faces did not (*M* = - .172 μV; *SD* = .389 μV).

**Figure 11.**
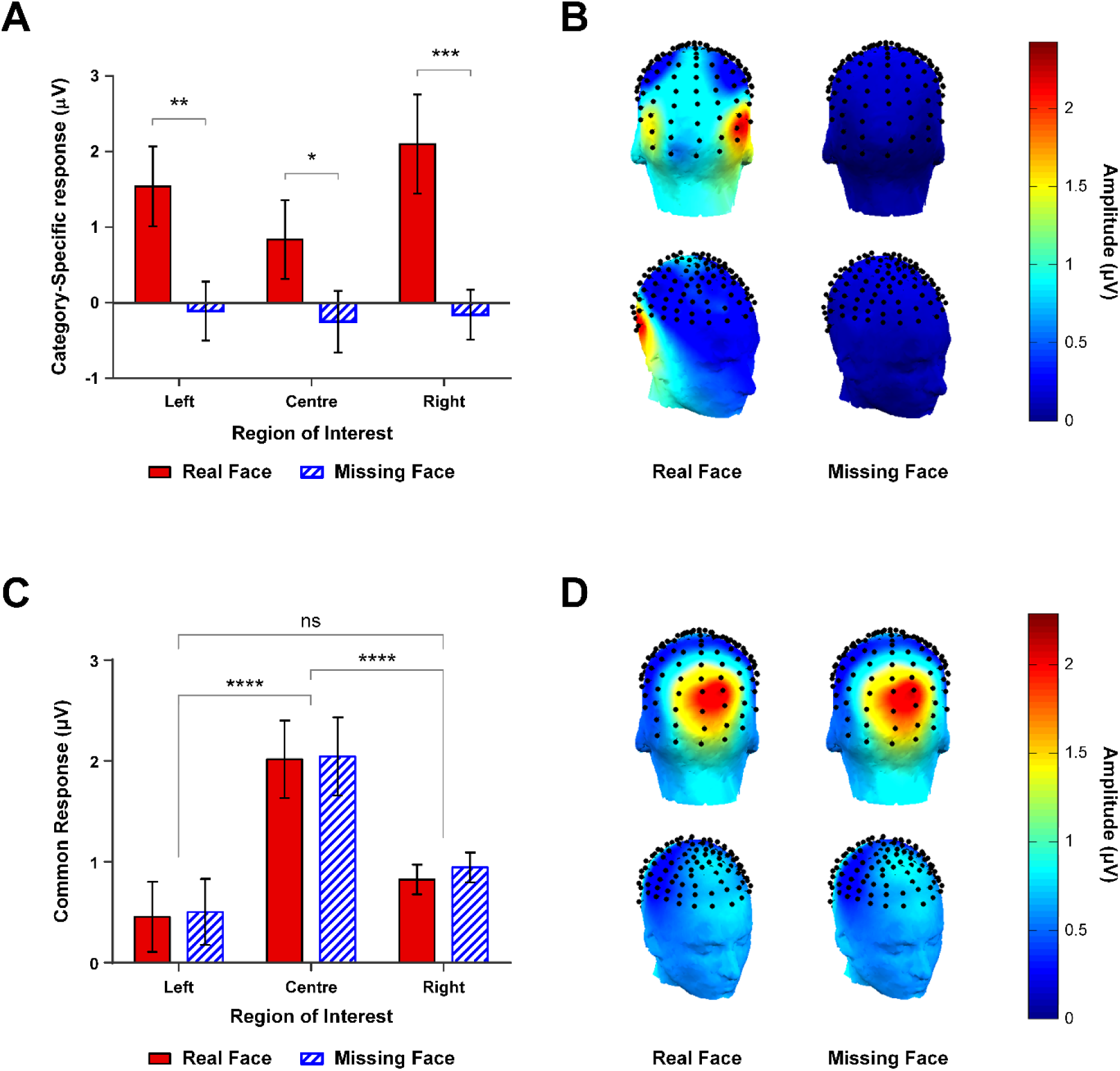
Sum of baseline-corrected amplitudes for the false-sequence data in Expt. 2. **(A)** The category-selective response (i.e., the sum of the first 8 harmonics of 1 Hz) was strongly modulated by both ROI and false-sequence type. In all ROIs, real faces (solid red) generated a clear differential response, where “missing faces” (blue stripes) did not. **(C)** In contrast, the common response (i.e., the sum of the first 4 harmonics of 12 Hz) did not differ with false-sequence type, but was strongly modulated by ROI. Significance codes: ^****^ p < .0001; ^***^ p < .001; ^**^ p < .01; ^*^ p < .05. Error bars are within-subjects 95% CIs, overlap should not be interpreted by eye (Cumming & Finch, 2005). **(B)** & **(D)** scalp topographies corresponding to the category-selective and common responses, shown for real and missing faces.

#### 7.3.2. False-sequence response at the common frequency (12 Hz).

Again pooling across all channels and conditions, we observed a significant response *(p* < .001, one tailed) at each of the first four harmonics of the common frequency (i.e., 12 Hz). At the level of ROI, a two-way within subjects ANOVA with the factors *Sequence Type* and *ROI* revealed that there was no interaction between these factors, *F*(2,20) = .876, p = .432. Moreover, the common response for the false-sequences was not modulated by *Sequence Type, F*(1,10) = 2.98, *p* = .115. As can be seen in Figure 11C, the magnitude of the common frequency response was comparable for missing face sequences (*M* = 1.16 μV; *SD* = 0.918 μV) and real face sequences (*M* = 1.10 μΥ; *SD* = 0.926 μV). In contrast, there was a strong main effect of ROI on the common frequency response, *F*(2,20) = 23.65, *p* < .0001. Follow-up Tukey contrasts showed that the common response for the central ROI (*M* = 2.03 μΥ; *SD* = .973 μV) was significantly larger than both the left ROI (*M* = .479 μΥ; *SD* = .270 μV), *z* = - 6.95, *p* < .0001, *d* = 1.509, and right ROI (*M* = .884 μV; *SD* = .464 μV), *z* = −5.14, *p* < .0001, *d* = 1.682. However, there was no difference between the two lateral ROIs, *z* = 1.82, *p* = .164, *d* = 0.817.

### 7.4. Time-Domain Results

Figure 12 shows the global response in the time-domain to missing and real faces, both before and after filtering out the common response at 12 Hz. Where the category-selective response for real faces was comprised of the same four spatiotemporal components as seen in Expt. 1, there was hardly any category-selective response for missing faces. That is, the response evoked by missing faces appeared to be almost entirely captured by the common response at 12 Hz.

**Figure 12.**
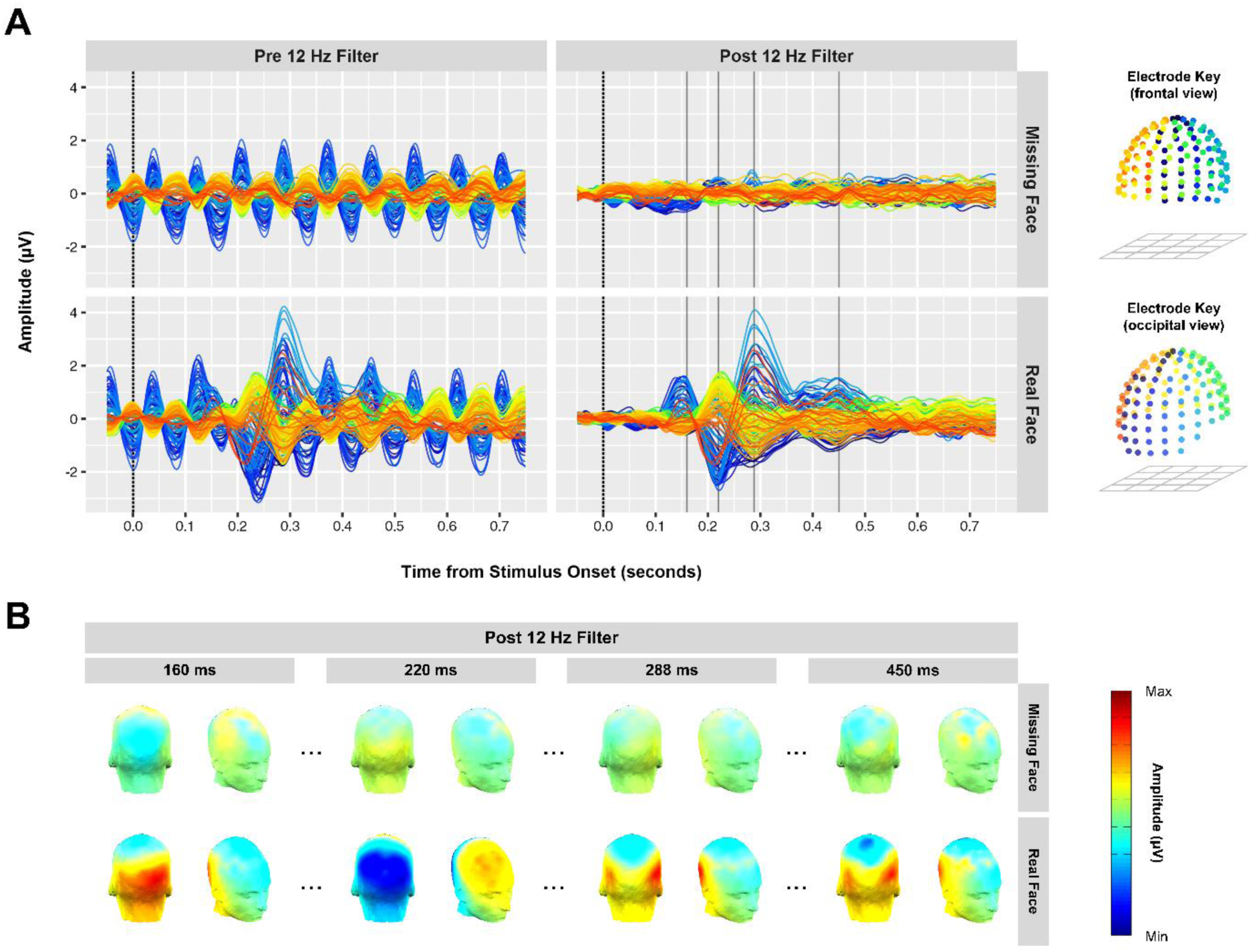
Time-domain response to missing faces and real faces in Expt. 2. **(A)** Conditional mean waveforms time-locked to stimulus onset for missing faces (top row) and real faces (bottom row). Left column: evoked response before applying 12 Hz notch filter to remove the response common to both faces and objects. Right column: The differential response to faces after notch-filtering at 12 Hz. Only real faces give rise to a clear differential (i.e., face-selective) response, comprised of the same four spatiotemporal components seen in Expt. 1. **(B)** Scalp topographies for missing and real faces at time points corresponding to the four components of the face response (160 ms, 220 ms, 288 ms, and 450 ms). Note that amplitude scales are fixed for condition pairs at each time point.

For each condition separately, we assessed the presence of a category-selective response in each ROI by comparing the response with a zero baseline in a series of paired t-tests. As can be seen in Figure 13, where the components evoked by real faces reached significance according to our imposed criteria, the response to missing faces did not approach either cut-off in any ROI. Although the averaged response of the four channels in the right ROI was not considered significantly different from a zero baseline according to either significance criteria, an inspection of Figure 12A suggests there might be a small increase on several occipito-temporal channels in the missing face condition around 300 ms. Given that this “blip” temporally coincides with the P3-face component, for interests’ sake we further inspected the response on individual channels. The missing face response on a single channel (PO12) met our less conservative significance criterion (i.e., *P* < .01, uncorrected) for a brief duration of 13.67 ms around 277 ms. On balance, these time-domain analyses suggest that where the real faces elicited a typical four component differential response (e.g., Retter and Rossion 2016), missing faces did not evoke an identifiable differential response. In other words, there was no evidence that objects replacing expected faces were processed differently to any other object in the rapid sequence.

**Figure 13.**
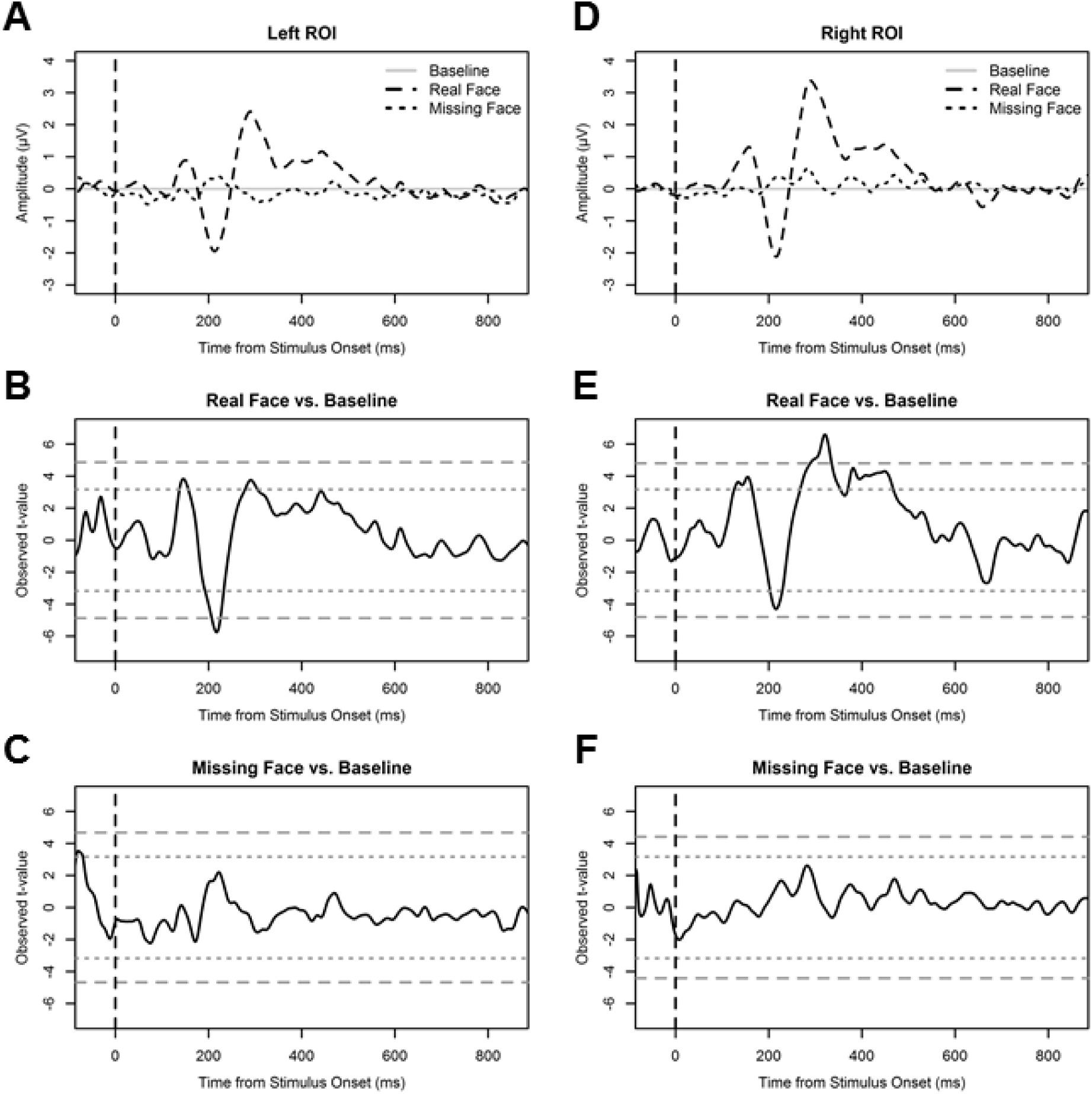
Time-domain analysis results in Expt. 2. Left Column: **(A)** Conditional mean amplitude in the left ROI following 12 Hz filtering, shown as a function of time from stimulus onset (vertical dashed line). **(B)** Left ROI analysis of the ‘real face’ response (real face vs. baseline), showing observed t-values for each time point. **(C)** Left ROI analysis of the missing face response (missing face vs. baseline), showing observed t-values for each time point. **(D)**, **(E),** & **(F)** are the same panels for the right ROI. Horizontal dashed lines are significance criteria (long dash = *p* < .05, permutation corrected; short dash = *p* < .01, uncorrected). Note that for both ROIs, only observed *t*-values in the real face condition met either criterion for significance, suggesting the response in the missing face condition did not differ from baseline at any point following stimulus onset.

## 8. Experiment 2 Interim Discussion

Our aim in Expt. 2 was to examine whether rare omissions of highly temporally predictable faces in an FPVS sequence would evoke an expectation related response. If this were the case, then the response to object images that replaced expected faces (so-called “missing faces”) should differ from the response evoked by other object images in the FPVS sequence. Our results here do not support this suggestion. Frequency-domain quantification of the differential response to both missing faces and real faces showed that where real faces elicited a strong category-selective response, the analogous response to missing faces was at floor level, not significantly greater than zero. Moreover, the response to missing faces in the time-domain was completely captured by the 12 Hz common response. This suggests that the response evoked by objects replacing expected faces was no different to that evoked by any other object in the sequence. Based on these results, we conclude that temporal expectation induced by the periodicity of critical stimuli in FPVS sequences is not sufficient to generate the category-selective response yielded by this paradigm.

As a secondary point, it is worth noting that even though the overall periodicity of the faces in these sequences was degraded (i.e., 10% of periodic faces replaced by objects), our initial frequency-domain analysis of the intact sequences still revealed a clear category-selective response. This is an encouraging finding, since it suggests the category-selective response measured with rapid continuous visual stimulation designs can tolerate a small amount of imperfection in the periodicity of critical category exemplars (note that the exact amount of degradation remains an empirical question). This advantage, along with the high signal-to-noise ratio offered by the technique, make FPVS an ideal method for testing participants who may particularly noisy EEG signals, or difficulty following instructions about tasks or blinks, such as young infants (see de Heering A and Rossion 2015), or psychiatric patients.

## 9. General Discussion

The goal of our study was to establish whether the category-selective signal elicited by the periodic appearance of critical category exemplars in a highly dynamic visual stimulation stream is generated or modulated by participants’ temporal expectations about those exemplars. To this end, we examined *i)* whether faces embedded in a continuous stream of objects at either periodic or nonperiodic intervals elicit different neural responses (Expt. 1), and *ii)* whether there is an identifiable neural response to a violation of the faces’ temporal predictability (Expt. 2).

The research we have reported here establishes several results. First, the category-selective response for faces embedded in a continuous stream of object images does not vary as a function of the faces’ temporal predictability. In Expt. 1, both frequency-domain and time-domain based analyses found no evidence that the category-selective response differs between faces embedded at periodic or nonperiodic intervals in the FPVS sequence. That is, the magnitude of the response and its spatiotemporal unfolding were entirely comparable for temporally predictable and unpredictable faces. That the response to periodic and nonperiodic faces was indistinguishable at the scalp level speaks directly to the concern that the category-selective response in FPVS may in fact be driven by the temporal predictability of the critical stimuli - our data emphatically suggest this is not the case. Importantly, since even the relatively late category-selective components were insensitive to temporal predictability (e.g., the P3-face, peaking around 300 ms), these results suggest that the entire differential response to faces in FPVS designs reflects a categorization response, rather than a response due to temporal expectancy. A second major finding is that there is no category-selective response for rare omissions of highly temporally predictable faces. In Expt. 2, we showed that the response evoked by object images that unexpectedly replace temporally predictable faces is no different to that evoked by other object images in the sequence - images about which participants have no temporal expectations. In other words, any temporal expectations induced by the periodicity of faces in our sequences was not sufficient to generate or modulate a category-selective response in the absence of a true face stimulus.

That temporal predictability plays no role in driving the category-selective signal in dynamic visual streams is somewhat surprising given the wealth of temporal expectation effects documented in the literature (Ariga and Yokosawa 2008; Mathewson et al. 2010; Kok, Jehee, et al. 2012; Rohenkohl et al. 2012; Cravo et al. 2013; Breska and Deouell 2014; Kok et al. 2014; Morillon et al. 2016). Here we consider a number of explanations for these conflicting findings. As suggested above, one possibility is that the periodicity of faces in our two experiments did elicit temporal expectations in participants, but that the high saliency of face stimuli for the visual system (Hershler and Hochstein 2005; Crouzet et al. 2010) evoked a very strong response that was simply too robust to be modulated by temporal expectation. On this possibility, if we had used a less potent critical stimulus category with this approach (e.g., pictures of houses or body parts, Jacques et al., 2016), or degraded/ambiguous face stimuli (Zhang et al. 2008), we might well have observed an effect of temporal expectation. While we cannot rule out this ‘robust-to-expectation-effects’ explanation on the basis of the current data (as the critical category was faces in both experiments), it is somewhat undermined by the fact that other forms of top-down expectation (e.g., category level expectation, such as expecting to see a face instead of a house), have been shown to be perfectly capable of modulating the neural response to face stimuli (Gregory 1970; Esterman and Yantis 2009; Puri et al. 2009; Egner et al. 2010; Jiang et al. 2015).

An alternative possibility is that the periodicity of faces in our FPVS sequences simply did not engender temporal expectations about faces in our participants. Although a number of studies have shown that rhythmic visual stimulation does indeed induce temporal expectations in observers (Ariga and Yokosawa 2008; Mathewson et al. 2010; Rohenkohl et al. 2012; Cravo et al. 2013; Breska and Deouell 2014), in our case the high visual processing load may have undermined participants’ ability to form reliable and accurate temporal expectations about the faces in the sequences. The FPVS approach places the visual system under significant strain - observers see a very large number of images (e.g., 720 images in a single minute) at an extremely high frequency of presentation (e.g., 12 images per second), with each image viewed for just 83.33 ms before being replaced. Within this high-load stimulation, the critical periodicity participants must entrain to is in fact an *embedded* periodicity (i.e., the face presentation frequency at 0.67 Hz, not the common frequency at 12 Hz). Where it is perhaps relatively easy for participants to perceive the rhythmic ‘beat’ (Rohenkohl et al. 2012) of the common frequency as each new image appears, it may be comparatively harder to perceive the beat of the face instances. Indeed, we are not aware of any study thus far that has examined temporal expectation effects for embedded visual periodicities. Moreover, we note that the face stimuli used here were not typical, full frontal face exemplars of uniform size, but rather varied widely in terms of viewing angle, face size, lighting, background, etc. As such, any temporal expectations about these faces would have been obliged to rely on a high-level, category-based stimulus template, rather than a template based on low-level commonalities between face exemplars. These temporal and image-based complexities distinguish our study from existing rhythmic visual stimulation studies, which have typically used slower presentation rates and/or shorter, less complex visual stimulations (Mathewson et al. 2010; Rohenkohl et al. 2012; Cravo et al. 2013; Breska and Deouell 2014). Where periodicity certainly engenders temporal expectations under these simplified conditions, we have shown here that it does not do so when visual stimulation is dynamic and continuous, simulating ecological conditions of natural vision.

Still another consideration is whether the impact of temporal predictability on stimulus processing might vary with the task-relevance of that stimulus. To the best of our knowledge, previous studies that have demonstrated modulation of perceptual processing by rhythmic temporal expectations have used behaviorally relevant target items (Ariga and Yokosawa 2008; Rohenkohl et al. 2012; Cravo et al. 2013; Breska and Deouell 2014; Morillon et al. 2016). That is, the stimulus for which an effect of temporal expectation was observed was one that participants actually responded to in some way. In contrast, the faces in our experiments were never task-relevant for participants, who we instructed to monitor the central fixation cross for color changes. Note that we are not suggesting that task-irrelevant stimuli cannot entrain temporal expectations in participants - indeed, several studies have already documented this (e.g., den Ouden et al. 2009; Alink et al. 2010) - but that perhaps in order for temporal expectations to facilitate perceptual processing, the critical stimuli must be attended to in some way (i.e., must be task-relevant). In the present experiments, we deliberately imposed an orthogonal task on the stimulation sequence, so as to disentangle the often confounded effects of attention and expectation (Summerfield and Egner 2009). However, if participants were required to judge some aspect of periodic faces explicitly (e.g., “respond when you see a female face”), we might well have observed an effect of temporal predictability on the category-selective response (Kok, Rahnev, et al. 2012).

Taken together, the findings reported here provide strong support for the claim that the index of perceptual categorization yielded by dynamic visual stimulation approaches (e.g., Rossion et al. 2015; Jonas and Rossion 2016; Retter and Rossion 2016) is immune to the temporal predictability of critical category stimuli. Importantly, this finding not only validates the use of FPVS as an objective tool with which to operationalize perceptual categorization processes, but also has important implications for understanding human perceptual categorization in a rapidly changing (i.e., dynamic) visual scene. The data reported here undermine a predictive coding interpretation of category-change detection in the human brain (Rao and Ballard 1999; Friston 2005; Alink et al. 2010; Kok, Jehee, et al. 2012), in that temporally predictable faces in our study were not associated with reduced neural activity (due to their ‘redundancy’), or a sharpened sensory representation (due to noise suppression) (Kok, Jehee, et al. 2012). That top-down temporal expectations do not facilitate sensory processing in the context of a dynamic continuous scene points to the interesting possibility that predictive mechanisms are not automatic, but rather subject to the rate at which visual stimulation changes. In this way, the research reported here highlights a broader need to test theoretical predictions under more ecologically relevant conditions, simulating the strain that natural vision in the real world places on our visual system.

### 9.1. Conclusion

This research establishes that category-selective neural responses elicited in a dynamic visual stream are immune to the temporal predictability of the critical category exemplars. As such, this category-selective response can be taken as a relatively pure index of perceptual categorization. We suggest that most likely explanation for why temporal expectation plays no role in driving the category-selective response is that high visual processing load in dynamic visual stream designs undermines observers’ ability to generate reliable temporal expectations. Additionally, the immunity of the category-selective response to temporal predictability may be tied to the use of an orthogonal task that renders the critical stimuli task-irrelevant.

## 10. Acknowledgements

The authors are grateful to T. Retter for her helpful insights and expertise in developing the false-sequencing data analysis. This work was supported in part by the European Research Council to BR [grant no: facessvep 284025] and cofounded by the University of Louvain and the Marie Curie Actions of the European Commission [grant no: F211800012] to GLQ.

## Footnotes

1 Note that unlike some previous studies (den Ouden HE *et al.* 2009; Kok P *et al.* 2014) in which a stimulus omission means there is no visual input at all, here the expected stimulus (a face) is *replaced* with an unexpected stimulus (an object).

2 For completeness’ sake we also inspected the common response magnitude in a central ROI identical to that used in Expt. 1 (i.e., averaged over POOz, Oz, POO6, PO4h), and found it to be just 0.0055 μν lower than that given by the central ROI we used in the main analyses (i.e., averaged over POOz, O2, POO6, PO4h)

3 Although the first three harmonics of 1 Hz did not reach our significance criterion, we had no a priori reason to exclude the signal on these harmonics from our quantification.

